# Brain Laterality Dynamics Support Human Cognition

**DOI:** 10.1101/2021.08.05.455350

**Authors:** Xinran Wu, Xiangzhen Kong, Deniz Vatansever, Zhaowen Liu, Kai Zhang, Barbara J Sahakian, Trevor W Robbins, Jianfeng Feng, Paul Thompson, Jie Zhang

## Abstract

Hemispheric lateralization constitutes a core architectural principle of human brain organization underlying cognition, often argued to represent a stable, trait-like feature. However, emerging evidence underlines the inherently dynamic nature of brain networks, in which time-resolved alterations in functional lateralization remain uncharted. Integrating dynamic network approaches with the concept of hemispheric laterality, we map the spatiotemporal architecture of whole-brain lateralization in a large sample of high-quality resting-state fMRI data (*N*=991, Human Connectome Project). We reveal distinct laterality dynamics across lower-order sensorimotor systems and higher-order associative networks. Specifically, we expose two aspects of the laterality dynamics: laterality fluctuations, defined as the standard deviation of laterality time series, and laterality reversal, referring to the number of zero-crossings in laterality time series. These two measures are associated with moderate and extreme changes in laterality over time, respectively. While laterality fluctuations depict positive association with language function and cognitive flexibility, laterality reversal shows a negative association with the same neurocognitive factors. These opposing interactions indicate a dynamic balance between intra- and inter-hemispheric communication, i.e., segregation and integration of information across hemispheres. Furthermore, in their time-resolved laterality index, the default-mode and language networks correlate negatively with visual/sensorimotor and attention networks, indicating flexible while parallel processing capabilities that are linked to better out-of-scanner cognitive performance. Finally, the laterality dynamics correlate with regional metabolism and structural connectivity and showed significant heritability. Our results provide insights into the adaptive nature of the lateralized brain and new perspectives for future studies of human cognition, genetics and brain disorders.

## Introduction

Hemispheric lateralization is a prominent feature of human brain organization ^1^, with inter-hemispheric differences repeatedly observed in both structure and function ^2–5^. For example, the left planum temporale, commonly referred to as the Wernicke’s area, shows reliable activity in cognitive paradigms that probe auditory processing and receptive language ^6,7^. In addition to such reports from “task-induced activation” studies on functional lateralization ^4,8^, emerging evidence also indicates lateralization within the human brain’s intrinsic connectivity architecture at rest. For example, recent neuroimaging studies suggest that hemispheric lateralization, estimated using network-based approaches on resting-state fMRI data, can accurately predict activity-based lateralization during cognitive task performance ^9,10^. Together, existing evidence alludes to the vital contribution of hemispheric lateralization to healthy and adaptive mentation.

Functional lateralization is traditionally considered as a static and trait-level characteristic of individuals ^5,9,11,12^ that is hypothesized to enhance neural capacity ^13,14^. For example, laterality of language and attention networks have been previously associated with individual differences in linguistic and visuospatial abilities ^5,9,11,12^. Beyond such trait-level characteristics, however, recent evidence also suggests the “dynamic” nature of intrinsic brain networks over time ^15–17^. Dynamic functional connectivity can track the time-varying alterations in cognitive states, task demands and performance ^18–20^, providing insights into how brain network reconfiguration relates to cognition, consciousness, and psychiatric disorders ^16,19,21,22^. In parallel, emerging findings now indicate that the degree of lateralization is instantaneously modulated by various external factors such as attention, task contexts and cognitive demands ^23–26^, which may arise from time-varying interactions between bottom-up and top-down neural processing ^4,27^. Therefore, it is possible that hemispheric lateralization also changes across time to accommodate changing demands of the environment, which may be evaluated by dynamic brain network approaches.

However, these two vital aspects of brain network interactions, in other words the dynamic changes of brain lateralization and their relationship to higher order cognition have not been explored to date. Therefore, in the present study we developed a measure of “dynamic lateralization” and tested its significance in explaining individual variability in cognitive performance. Specifically, we investigated the laterality dynamics by two complementary measures, i.e., laterality fluctuations and laterality reversal, which reflect moderate and extreme changes in laterality, respectively, on intrinsic brain networks constructed from high-quality resting-state fMRI data from the Human Connectome Project (HCP) ^28^. Here, we show that laterality fluctuations are positively associated with language function and cognitive flexibility, while laterality reversal shows an opposite effect, suggesting a balance between intra- and inter-hemispheric information communication. Furthermore, negative correlations in time-varying laterality between default-mode network and visual/sensorimotor and attention networks and their relationship with cognitive performance were revealed, suggesting parallel information processing capacity which may facilitate adaptive cognition. Additionally, we also investigated the neural and anatomical factors that may affect dynamic laterality of the human brains and established the heritability of the dynamic laterality measures.

## Results

### Dynamic Laterality Index

We analyzed resting-state fMRI data from 991 subjects in the HCP cohort, and extracted BOLD time series of all 360 cortical regions using a group-level parcellation scheme (HCP MMP1.0) ^29^. To map the time-varying lateralization architecture, we developed a measure termed dynamic laterality index (DLI) by calculating the laterality index in each sliding window for each region of interest (ROI) (Fig. 1a). Specifically, we adopted a global-signal based laterality index that can effectively capture brain lateralization characteristics underlying higher order cognition ^11^, defined as the difference between a ROI’s BOLD correlation with the global signal of left brain and its correlation with the right brain at each time window (see Methods). DLI of a ROI (a time series of laterality index) is then obtained by calculating the laterality index for each time window, see Fig. 1a. Positive DLI of a region indicates stronger interaction with the left hemisphere (i.e., leftward laterality), while a negative one indicates rightward laterality.

**Figure 1.**
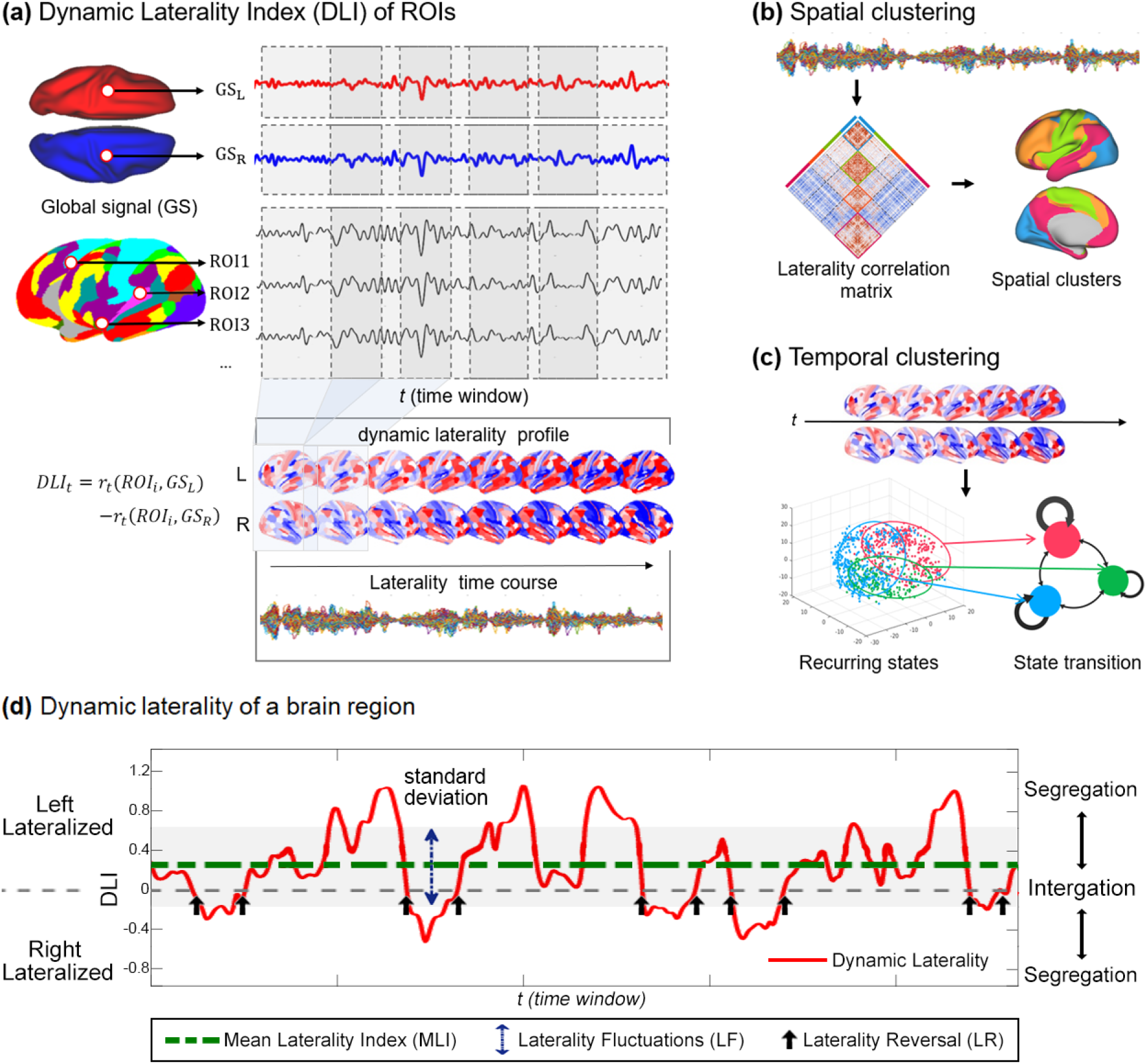
The workflow of dynamic laterality analysis. (a) Definition of the dynamic laterality index (DLI). The DLI of ROI_i_ within time window *t* is defined as the correlation coefficient (z-transformed) between Global Signal of the left hemisphere (*GS_L_*) and the ROI minus the correlation coefficient betwee) and the ROI minus the correlation coefficient between the Global Signal (*GS_R_*) of *the* right hemisphere and the ROI. Using a sliding window approach, we obtained a time series of dynamic laterality index for each ROI. (b) Laterality correlation matrix, which is obtained by correlating laterality time series across all ROIs. Spatial clustering is then performed to identify spatial clusters of brain regions demonstrating covariation in laterality time series. (c) Temporal clustering of whole-brain laterality patterns, which identifies potential recurring laterality patterns. (d) Illustration of dynamic laterality index and relevant dynamic laterality measures using a ROI that is left-lateralized. The red curve represents the time series of dynamic laterality index. The green dotted line is the mean laterality index (MLI, 0.26), and the blue double arrow denotes the standard deviation of the laterality time series, which measures the level of laterality fluctuations (LF). The black arrow represents the laterality reversal (LR, the change of the sign of lateralization across two consecutive time windows). Large magnitude of laterality index indicates segregation at the hemispheric level, while small magnitude of laterality index (near 0) indicates integration across two hemispheres.

We demonstrate that the global-signal based laterality index we adopted is highly correlated to a conventional laterality measure Autonomy Index ^9,30^ that is widely used, but has many additional advantages. Autonomy Index is defined as the difference between the overall functional connectivity of a ROI with the left hemisphere and its overall functional connectivity to the right hemisphere, see Methods. We show mathematically that these two laterality index are highly correlated (Supplement methods). Using resting-state fMRI data from HCP, we found high correlation coefficient between the whole-brain laterality profile obtained by the mean dynamic laterality index (MLI, average across all time windows) and that obtained by the traditional Autonomy Index (r= 0.64±0.12, all subjects have p<0.05, Supplemental Fig.1). The advantage of the global-signal based laterality index is its higher robustness across different scans. The mean laterality index has better replicability over 4 sessions involving left- and right-scans (correlation across sessions: 0.66±0.018) compared to Autonomy Index (0.41±0.10), see Supplemental Fig. 1. Furthermore, global-signal based laterality index is efficient in that it only takes 2*n time, compared to the traditional Autonomy Index that takes n^2^ time (n being the number of regions), therefore is economic for multiple time windows. Considering its validity, replicability and computational advantages, we used this global-signal based laterality index in calculating dynamic laterality index. Finally, window length remains an important parameter in dynamic analysis. Our main results were based on a window length of 30 TRs (30*0.72=21.6s) and a step size of 1 TR. We confirmed all results with different window sizes (60 TRs and 90 TRs per window, see Supplementary Fig. 1c).

Using this dynamic laterality index, we characterized the time-averaged laterality of large-scale brain networks in resting-state. The left and right hemispheres largely illustrated positive and negative mean laterality, respectively (Fig. 2a). In the left hemisphere, regions in the language (MLI=0.17±0.08) and default-mode (MLI=0.14±0.07) networks showed strong leftward laterality [networks defined by Cole-Anticevic Brain-wide Network Partition ^31^]. In the right hemisphere, the cingulo-opercular (MLI=−0.11±0.07), dorsal attention (MLI=−0.10±0.06) and visual networks (MLI=−0.08±0.08, Fig. 2a) showed strong rightward laterality. Frontoparietal network illustrated strong laterality within both hemispheres (left: MLI=0.14±0.07, right: MLI=−0.15±0.07). Comparatively, bilateral sensorimotor regions (left: MLI=0.036±0.04, right: MLI=−0.022±0.05) and left visual areas (MLI=−0.025±0.06) depicted relatively weak mean laterality.

**Figure 2.**
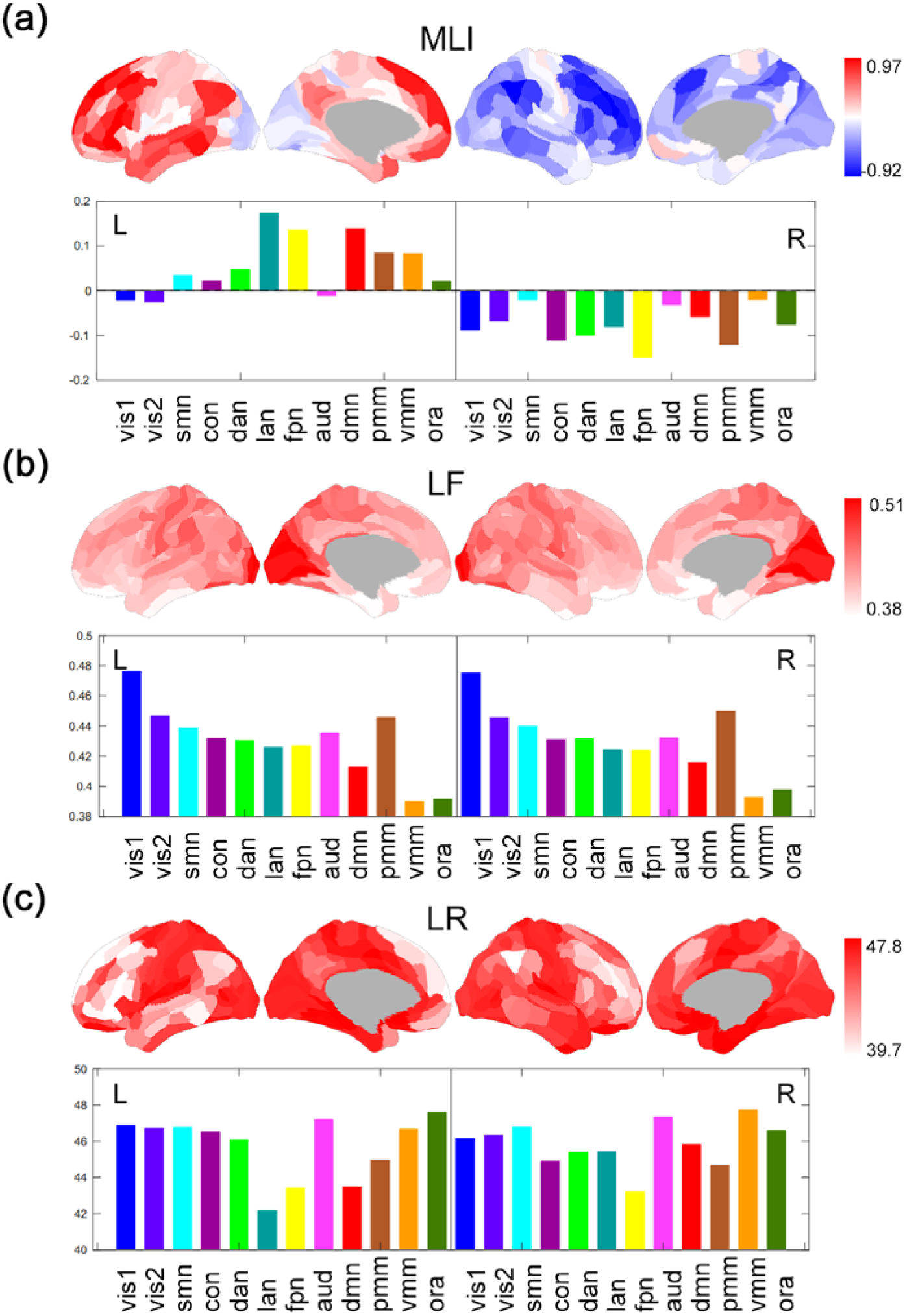
Dynamic architecture of whole-brain functional lateralization. (a) mean laterality index (MLI), i.e., time average of dynamic laterality index (b) Laterality fluctuations (LF) i.e., standard deviation of laterality time series, and (c) Laterality reversal (LR), i.e., the number of switches in the sign of laterality (from positive to negative or vice versa) of all 360 brain regions (averaged across 991 subjects), which are rearranged into 12 subnetworks by the Cole-Anticevic Brain-wide Network Partition. Vis1, Visual1; vis2, Visual2; smn, Somatomotor; con, Cingulo-Opercular network; dan, Dorsal-Attention network; lan, Language network; fpn, Frontoparietal network; aud, Auditory network; dmn, Default Mode network; pmm, Posterior-Multimodal; vmm, Ventral-Multimodal; ora, Orbito-Affective. L, left hemisphere; R, right hemisphere.

We further characterized the dynamic changes in laterality of a region from two different perspectives: the magnitude and the sign of laterality, by *laterality fluctuations* (*LF*) and *laterality reversal* (*LR*), respectively, see Fig. 1d. Laterality fluctuations is defined as the standard deviation of laterality time series, while laterality reversal specifically refers to the number of zero-crossings of laterality (switch between left- and right-laterality) in two consecutive windows. We illustrate laterality time series and these two measures using a brain region with high level of mean laterality (0.26, left-laterality, Fig. 1d). The standard deviation of time-varying laterality index corresponds to moderate changes, or deviations, with respect to the mean laterality. By contrast, laterality reversal reflects larger changes in laterality, corresponding to sufficiently extreme deviations from the mean, that results in sign-changes in laterality. Moreover, a large magnitude of the laterality index corresponds to segregation at the level of hemispheres, and a small magnitude (near 0) indicates integration across two hemispheres, see Fig. 1d and “network and structural basis of laterality dynamics” (Results) for explanation.

Based on the above two dynamic laterality characteristics, we found that the laterality of most brain regions varied considerably over time. Specifically, primary visual (LF=0.46±0.09; LR=46.8±1.7) and sensorimotor regions (LF=0.44±0.09; LR=46.6±1.9) generally illustrated stronger levels of variation in laterality (both laterality fluctuations and reversal) than higher-order association regions - including the frontoparietal (LF=0.43±0.11; LR=43.4±1.97), language (LF=0.43±0.10; LR=43.5±2.1), and default mode networks (LF=0.41±0.10; LR=44.7±1.7), see Fig. 2b and Fig. 2c.

### Spatial clustering of the laterality dynamics

After characterizing the temporal laterality fluctuations across the whole-brain, we investigated how spatially distributed regions that support different functional specializations correlate with each other in laterality (Fig. 1b). Through spatial clustering of the laterality time series across all brain regions, we identified four major clusters (Fig. 3a): Cluster 1 consisted mainly of regions from the bilateral visual network; Cluster 2 consisted of regions from the bilateral sensorimotor and cingulo-opercular networks; Cluster 3 mainly covered the frontoparietal network (more from the right hemisphere) and attention network; and Cluster 4 mainly consisted of regions from bilateral default-mode network, part of the frontoparietal network (mainly the left hemisphere) and regions from the language network (Fig. 3a). Cluster 1 and 2 showed higher levels of laterality fluctuations (reflected by higher standard deviation of dynamic laterality index and laterality reversal) when compared to Clusters 3 and 4 (repeated-measures ANOVA, p<0.001, pairwise t-test with p<0.01, see supplemental Fig. 3a).

**Figure 3.**
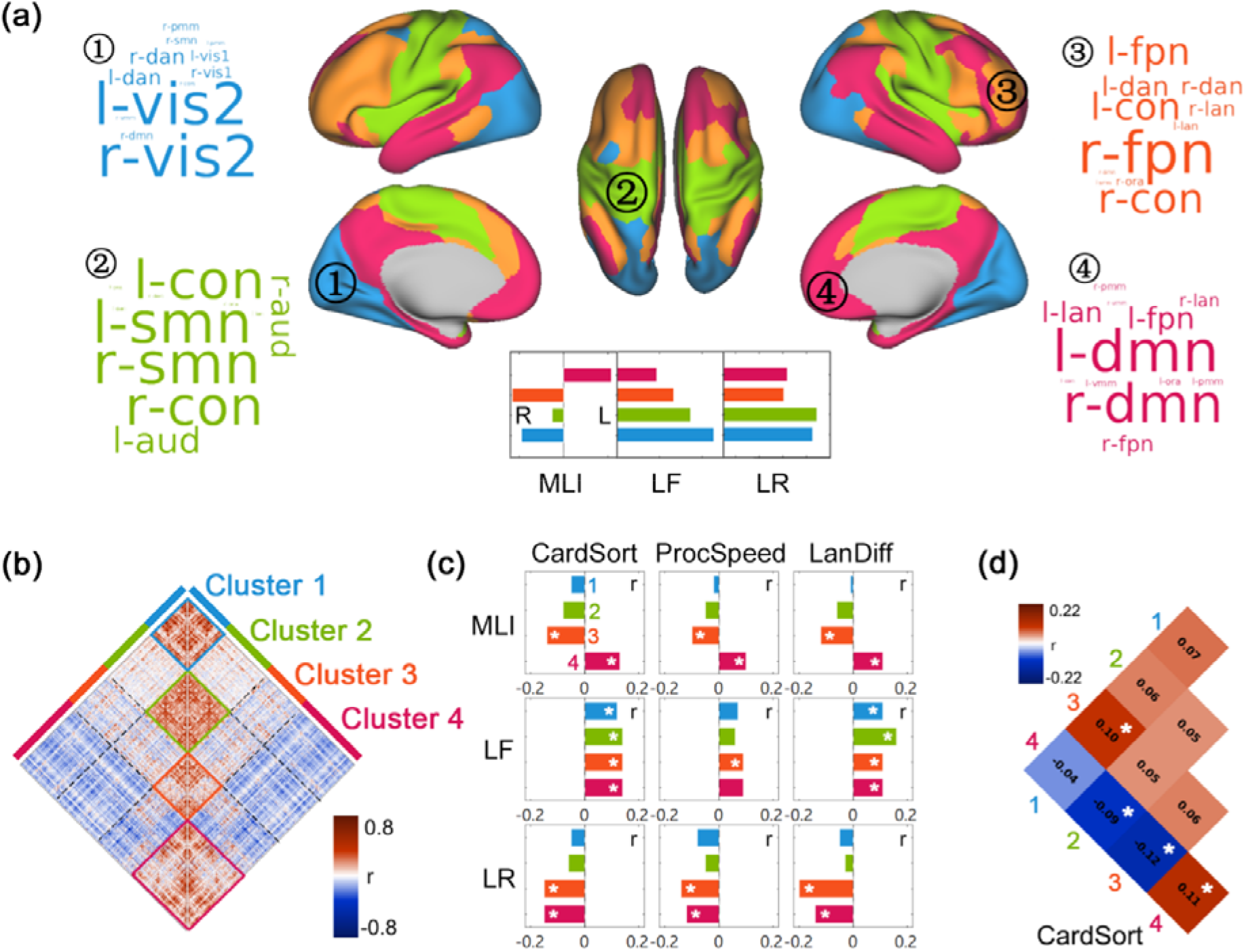
Spatial clustering of laterality dynamics. (a) Four spatial clusters revealed by clustering of the laterality time series of all 360 brain regions. The word-cloud for each cluster indicates the functional networks being involved (based on the Cole-Anticevic Brain-wide Network Partition), with the font size representing the proportion of each network in the cluster. L, left-lateralized; R, right-lateralized; MLI, Mean laterality index; LF, laterality fluctuations; LR, laterality reversal. The mean value of the three dynamic laterality measures of the four spatial clusters are shown in the inset. (b) The laterality correlation matrix across 360 brain regions (averaged over all 991 subjects). (c) The correlation between the three dynamic laterality measures (MLI, LF, LR) of the four spatial clusters and cognitive performance of three tasks. CardSort, the performance of Dimensional Change Card Sort Test; ProcSpeed, the performance of Pattern Comparison Processing Speed Test; LanDiff, mean difficulty of stories for each subject in HCP language task. (d) The association between the laterality correlation within/between clusters and the processing speed (Pattern Comparison Processing Speed Test). Only significant results (FDR corrected) are shown; see Supplementary Fig. 3 for all results.

We then explored the association between laterality dynamics and out-of-scanner cognitive performance. Specifically, we used multiple tasks adopted by HCP that are generally associated with either lateralized (e.g., language: story comprehension task, and attention: Flanker task) or bilateral (e.g., cognitive flexibility: Card Sort task, and working memory: List Sorting task) brain function. First, we found that the laterality fluctuations and laterality reversal of the identified spatial clusters correlated in a positive and negative manner, respectively, with cognitive performance. Second, significant correlation was only found for language function (story comprehension tasks), cognitive flexibility (Card Sort Test and pattern comparison task) and processing speed (Pattern Comparison Test) out of all 13 cognitive measures, see Supplemental Table 1.

Specifically, laterality fluctuations of Clusters 3 (FPN) and 4 (DMN and language network) correlated positively with the difficulty of stories a subject could understand (language task difficulty, LanDiff, Cluster 4: *r*=0.11, *p* =0.018; Cluster 3: *t*=0.11, *p*=0.014, Fig. 3c). A cognitive flexibility measure (Dimensional Change Card Sort Test, CardSort) also correlated positively with laterality fluctuations across all clusters (Cluster 1, r=0.12, *p*=0.001; Cluster 2, r=0.14, *p*<0.001; Cluster 3, r=0.14, *p*<0.001; Cluster 4, r=0.14, *p*<0.001). In contrast, laterality reversal of these clusters correlated negatively with cognitive performance: for language task difficulty, *r*=−0.2 (*p*<0.001) for Clusters 3, and *r* =−0.14 (*p*=0.002) for Clusters 4. For Card Sort task, *r*=−0.15 (*p*<0.001) for Cluster 3 and *r*=−0.15 (*p*<0.001) for Cluster 4. In addition, the performance of Pattern Comparison Processing Speed Test (ProcSpeed, the speed of completing the task) also showed positive association with laterality fluctuations of Cluster 3 (r=0.09, p=0.009) and negative association with laterality reversal of Cluster 3 (r=−0.14, p=0.001) and 4 (r=−0.12, p=0.003).

We furthermore explored how time-varying laterality time series of different clusters correlate with each other. Importantly, Cluster 4 showed negative laterality correlations with all other 3 clusters (Cluster 1: t=−81.7; Cluster 2: t=−84.37; Cluster 3: t=−62.1, all p<0.001, Fig. 3b), while Cluster 1, 2 and 3 showed positive correlation among themselves (Cluster 1 and 2, t=20.4; p<0.001; Cluster 2 and 3, t=15.2; p<0.001), indicating that Cluster 4 shows a tendency to lateralize in the hemisphere opposite to those of the other 3 clusters over time. The negative laterality correlation between Cluster 4 and Cluster 1, 2, and 3 also bear functional significance. We found that individuals with higher Card Sort score showed stronger negative correlation in laterality between Cluster 4 and Cluster 2 (r=−0.09, p=0.012), and Cluster 4 between Cluster 3 (r=−0.12, p=0.001), see Fig. 3d and Supplemental Table 2.

### Temporal clustering of laterality dynamics

We further explored the temporal organization of laterality dynamics by clustering whole-brain laterality state of multiple time windows (see Fig.1c and Methods). Three recurring laterality states (Fig. 4a) were identified, each showing distinct laterality patterns, different dwelling times and transition probabilities (Fig. 4b). Specifically, State 1 showed a typical leftward laterality in Cluster 4 (e.g., the left default-mode and language network; *t* =77.9, *p*<1e-20) and a rightward laterality in Cluster 2 (e.g., the sensorimotor network and the cingulo-opercular network; *t* =−109.3, *p*<1e-20) (Fig. 4a–4b). State 1 is the primary laterality state with the largest fraction of 41% and a mean dwelling time of 49.9±3.58 windows (about 35 second). State 2 showed a typical rightward laterality in Cluster 1 (the visual network; *t* =−108, *p* <1e-20), and a leftward laterality in Cluster 2 (the sensorimotor and the cingulo-opercular network; *t* =60.1, *p*<1e-20) and Cluster 4 (the language network; *t* =53.2, *p*<1e-20) (Fig. 4a–4b). State 3 showed a leftward laterality in Cluster 1 and 2 (the visual and sensorimotor network), and rightward laterality of Cluster 4 (e.g., the right default mode network, *t* =−64.7, *p* <1e-20) and Cluster 3 (e.g., the frontoparietal network in the right hemisphere, *t* =−41.4, *p* <1e-20). These two states showed similar fractions (State 2: 28%; State 3: 31%) and dwelling time (State 2: 10.5±2s; State 3: 12.1±2.3s).

**Figure 4.**
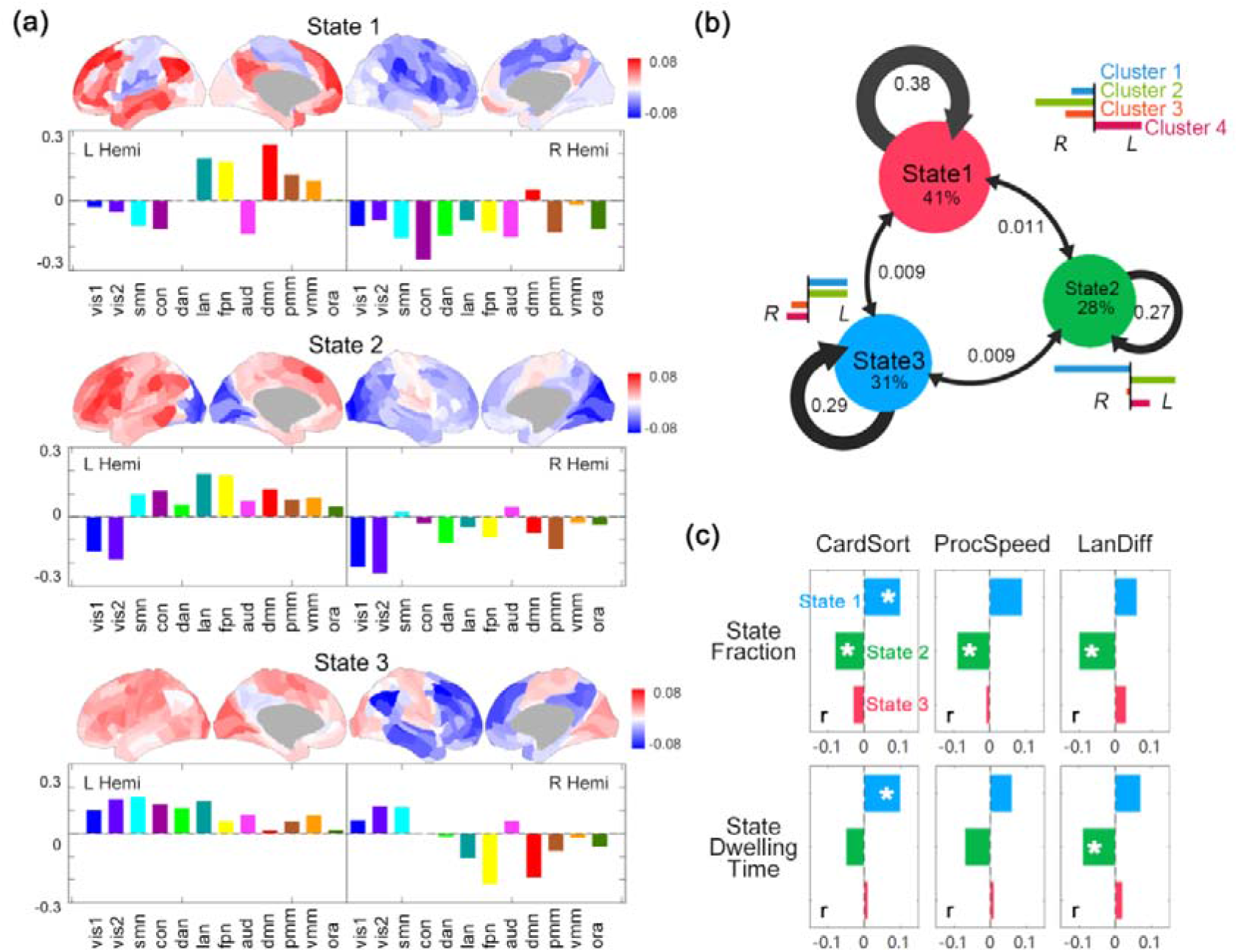
Temporal clustering of laterality dynamics. (a) The three recurring laterality states obtained by temporal clustering of the time-varying, whole-brain laterality states (360 regions) with colors indicating the network arrangement of the Cole-Anticevic Brain-wide Network Partition (CAB-NP). L, left hemisphere; R, right hemisphere. (b) The mean fraction of three states and the mean probability of switching between them. The bar plot next to each state represents the averaged laterality of the 4 spatial clusters in each state. L, left-lateralized; R, right-lateralized. (c) The correlation between the fraction/mean dwelling time of three states and memory ability (mem)/language ability. (d) The correlation between the probability of switching between/within the 3 states and language ability.

We further investigated the behavioral correlates of individual variations in the state transition properties. Results showed a link between State 2 and cognitive functions: individuals with less State 2 showed higher language task difficulty (fraction, *r*=−0.1, *p*=0.007; dwelling time, *r*=−0.09, *p*=0.015), higher CardSort performance (fraction, *r*=−0.08, *p*=0.014) and higher ProcSpeed score (fraction, r=−0.09, *p*=0.01). In addition, CardSort was positively correlated with the fraction (*r*=0.1, *p*=0.006) and dwelling time (*r*=0.1, *p*=0.014) of State 1. See Fig. 4c–4d and Supplemental Table 3.

### Network and structural basis of laterality dynamics

Next, we investigated how dynamic laterality is related to functional and structural brain network properties. First, we investigated how dynamic laterality of a brain region is related to time-varying network feature (including degree, participation coefficient, and modularity, reflecting large-scale organization of the brain like integration/segregation) and BOLD activity (amplitude of low-frequency fluctuation, ALFF). At the global level, we found that the whole-brain lateralization (the mean absolute value of DLI across all 360 regions) correlated negatively with the averaged degree (r=−0.43±0.1), participation coefficient (r=−0.2±0.12), interhemispheric connection (r=−0.52±0.11) and ALFF (r=−0.22±0.12), while showing positive correlations with modularity of the brain network (r=0.47±0.06) across time windows, see Supplemental Fig 4. This indicates that when the whole brain is highly segregated (modular), it also demonstrates high laterality; while an integrated state of the whole brain (low modularity) are accompanied by low laterality. At the local level, we found that these correlation patterns are most prominent for the high-order brain regions (most significantly in default-mode network, the language network and the frontoparietal network, Supplemental Fig. 8).

Second, we explored the structural correlates of laterality dynamics using diffusion MRI data. We found significant positive correlations between the laterality correlation matrix and structural connectivity matrix, including FA matrix (Spearman’s *rho*=0.14±0.02, significant in 99% subjects) and FN matrix (Spearman’s *rho*=0.14±0.02, significant in 99% subjects), see Fig. 5a. In line with this, homotopic regions generally showed positive correlations in their laterality time series (mean r=0.39∼0.79, Supplemental Fig. 2) due to the critical role of corpus callosum ^32^. These results suggested that brain regions with similar laterality dynamics tend to have stronger structural connection. Furthermore, laterality fluctuations (LF) of a region correlated positively with its FA/FN-degree, i.e., the sum of fractional anisotropy/fiber number of fibers between this region and all other regions (LF-FN: *rho*=0.32±0.09, significant in 99% subjects; LF-FA: *rho*=0.28±0.08, significant in 99% subjects, see Fig. 5b).

**Figure 5.**
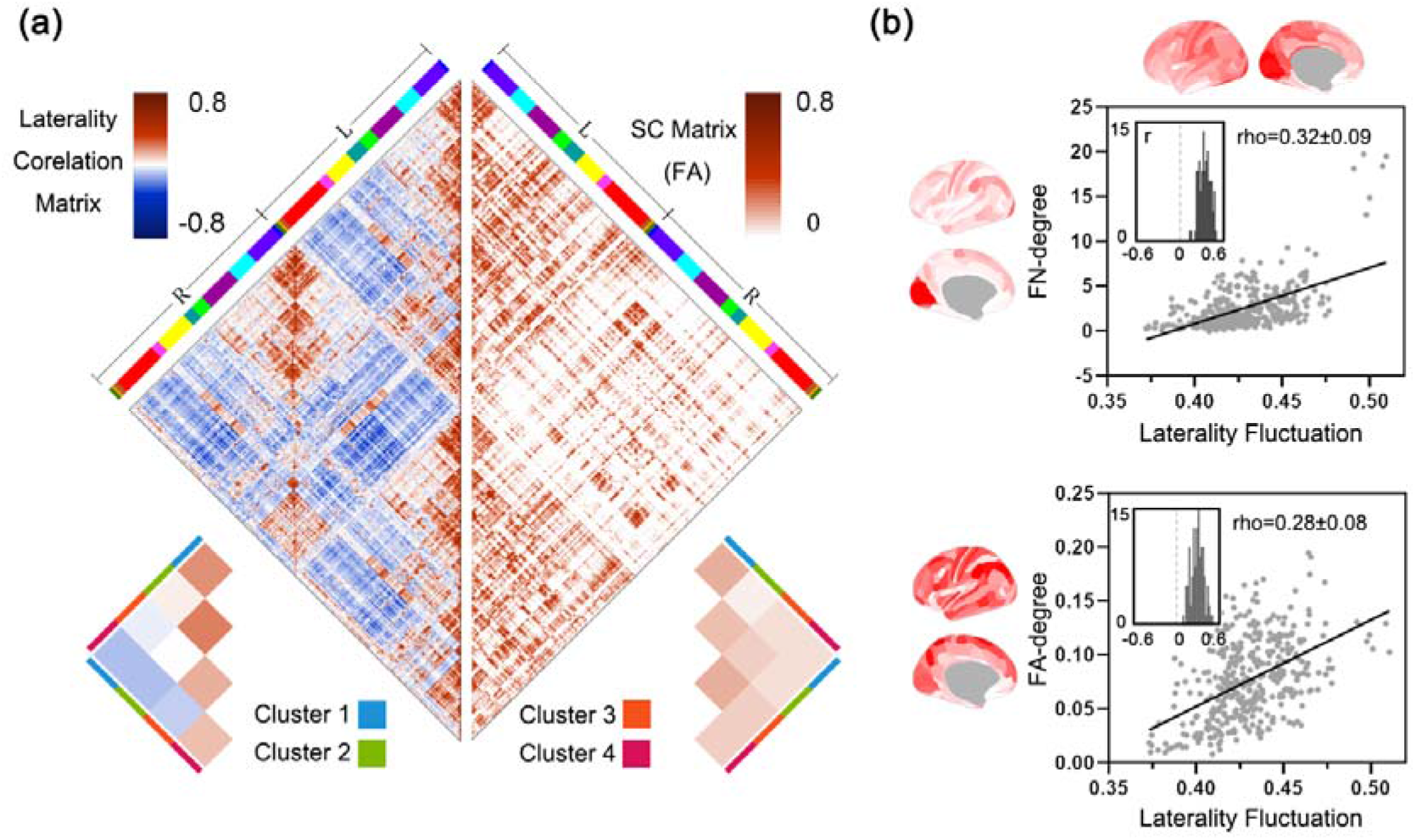
Structural basis of dynamic laterality. (a) Laterality correlation matrix (left half-matrix) and structural connection matrix (right half-matrix) demonstrates high level of correlation. These two matrices are obtained by averaging across all 99 subjects. L, left hemisphere; R, right hemisphere. The color ribbons indicate the network arrangement of CAB-NP detailed in Fig. 2. (b) Upper: The relationship between a structural measure (FN-degree) and the laterality fluctuations (LF) across all brain regions. Lower: The relationship between another structural measure (FA-degree) and the laterality fluctuations. The large scatter maps show the association between the averaged map of dynamic laterality measures and the averaged FA- /FN- degree map (both across all 99 subjects), and the inset shows the distribution of the individual correlation coefficients between the FA-/FN- degree map and laterality fluctuation map of each subject.

### Heritability of laterality dynamics

Finally, we investigated heritability of lateralization dynamics by twin analyses (using the kinship information). First we estimated heritability through the ACE model [additive heritability (A), common (C) and specific (E) environmental factors model]. We found that laterality fluctuations showed the highest heritability (*h*^2^) among all dynamic laterality measures (*h*^2^=0.09-0.48 for LF; *h*^2^ =0.002-0.41 for MLI; *h*^2^ =0.001-0.31 for LR, see Fig. 6a–c). Of all four spatial clusters, Cluster 4 (e.g., the default-mode network, part of the right frontoparietal network, and the language network) showed the highest heritability (MLI *h*^2^=0.19; LF *h*^2^=0.4; LR *h*^2^=0.06), followed by Cluster 3 (MLI *h*^2^=0.19; LF *h*^2^=0.36; LR *h*^2^=0.06. The heritability of Cluster 2 (MLI *h*^2^=0.13; LF *h*^2^=0.35; LR *h*^2^=0.03) and Cluster 1 (MLI *h*^2^=0.12; LF *h*^2^=0.33; LR *h*^2^=0.03) were relatively low.

**Figure 6.**
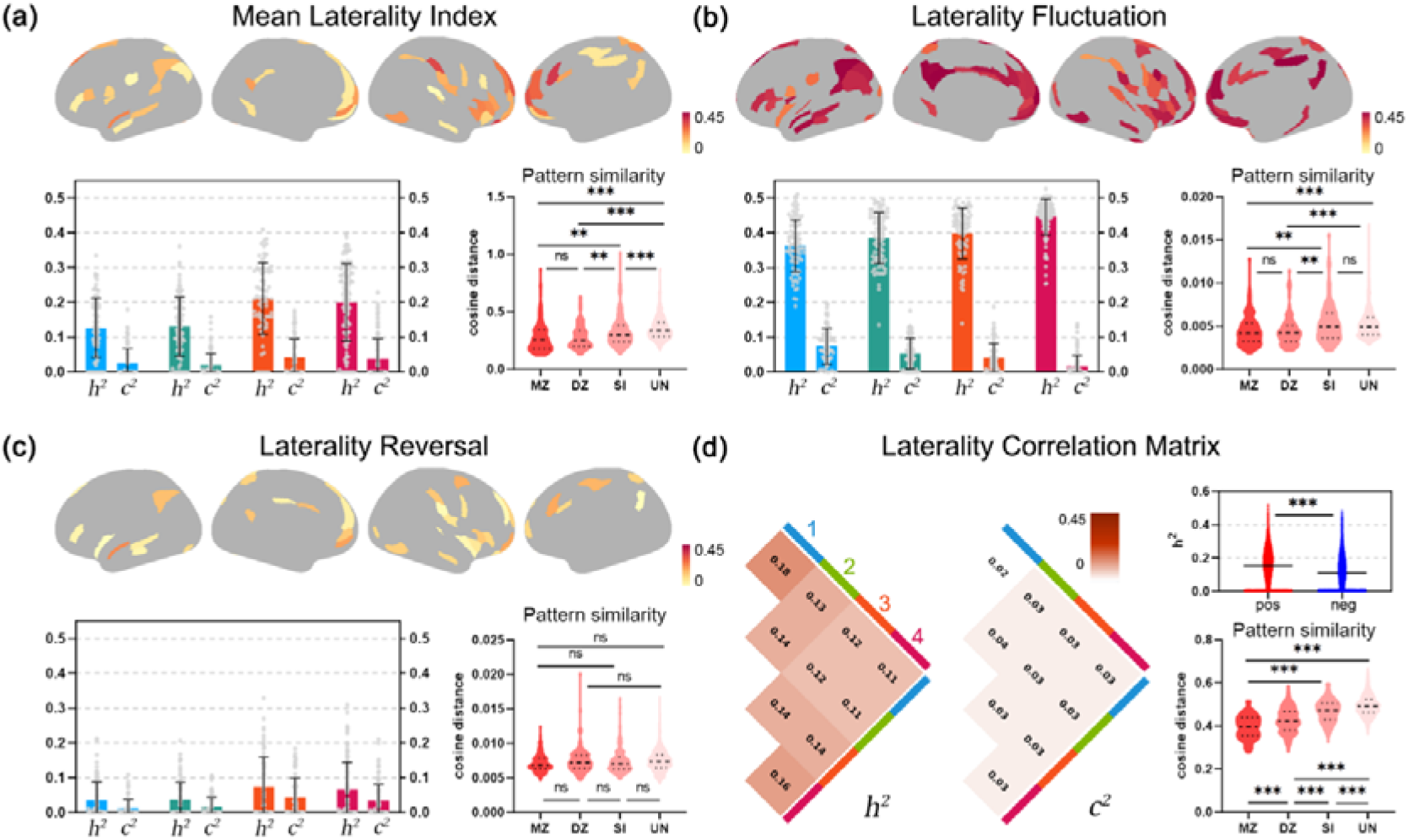
Heritability of dynamic laterality. (a-c) Heritability analysis of MLI, LF and LR. Upper. Heritability (*h*^2^) of each brain region. Only the regions with *h*^2^ of p<0.05 are retained. Lower Left: The *h*^2^ and common environmental factors (*c*^2^) for each cluster, and each data point represents a brain region. Lower Right: The cosine distance between whole brain map for each pair monozygotic twins (MZ), dizygotic twins (DZ), sibling (SI) and unrelated group (UN). Asterisk indicates the significant difference between the groups (one-way ANOVA). (d) Heritability of lateralization correlation matrix. Left: Average *h*^2^ and *c*^2^ in four clusters. Upper right: Differences in heritability between the positive and the negative links, i.e., links with positive or negative correlation in laterality. Lower right: The cosine distance of laterality correlation matrix of each pair of MZ, DZ, SI and UN. The error bar represents the mean value ±1 SD. One asterisk, p<0.05; two asterisks, p<0.01; three asterisks, p<0.001. Pos, positive correlation; neg, negative correlation.

We then calculated the cosine distance between each pair of monozygotic twins (MZ), dizygotic twins (DZ), sibling and unrelated group subjects using various dynamic laterality measures across the whole brain. As expected, the laterality correlation matrix showed greater similarity within MZ twins than those within DZ or non-twins (one-way ANOVA, *F*=152.7, *p*<0.0001; Fig. 6d). Similarly, the similarity of MLI and LF (of all regions) in twins, siblings and unrelated subjects, showed a decreasing trend, although the difference between MZ and DZ was not significant (for MLI, *F*=23.8, *p*<0.0001, MZ vs. DZ, *p*>0.99; for LF, *F*=10.8, *p*<0.0001, MZ vs. DZ, *p*=0.99; Fig. 6a–6b). There was no significant difference in LR between the four groups (*F*=1.48, *p*=0.22, Fig. 6c). In summary, dynamic laterality measures were generally heritable.

## Discussion

Laterality is traditionally argued to represent a stable, trait-like feature of the human brain. However, considerable amount of work has recently been devoted to characterizing dynamic functional connectivity (FC) that has been shown to be related to mental flexibility ^16^ and to predict ongoing mental states or task performance ^18^, which inspires the question of whether laterality of the brain is also fluctuating over time. The current study revealed the dynamic nature of laterality across the brain and its cognitive significance. Different levels of temporal variation of laterality were found across the brain: regions within the visual and sensorimotor networks generally showed higher variations, while regions from the default-mode, frontoparietal and language network showed relatively weaker variations. This suggested laterality in lower-order regions is more flexible possibly due to bilateral sensory inputs. Note this difference between lower-order and higher-order networks is not driven by dynamic FC, which demonstrates a reverse trend across lower and higher-order regions, i.e., sensorimotor and visual regions changes little in time for dynamic FC, while multimodal association regions change frequently ^33,34^. Dynamic FC characterizes micro-scale changes in brain networks while dynamic laterality reflects macro-scale, specifically, hemispheric-level changes.

To systematically characterize dynamic laterality changes, we used two measures, i.e., laterality fluctuations and laterality reversal, which showed positive and negative correlations with cognitive performance, respectively. These opposing associations suggest that these two measures capture different aspects of dynamic laterality. Generally, brain regions in the left and right hemisphere illustrated positive and negative mean laterality, respectively, (indicating they have more ipsilateral connections), with their laterality fluctuating around the mean value (Fig. 1). Greater fluctuations of laterality suggested larger number of states of intra- and inter-hemisphere interactions for a brain region. For example, in story comprehension tasks adopted, reading sentences with literal meanings has been shown to elicit activations in the left language areas (high left-laterality), while dealing with difficult metaphors were shown to involve the right homotopic regions (low left-laterality or even high right-laterality) as a compensation ^35^. Therefore, larger fluctuations of laterality are able to accommodate both specialized processing in one hemisphere (high left-laterality) and bilateral processing (low left-laterality or even high right-laterality) that is potentially beneficial in a difficult story comprehension task. This result is also consistent with findings indicating that the right hemisphere also makes significant contributions to language processing ^36,37^. For cognitive flexibility (Card Sort task) that generally involves both hemispheres ^38^ and larger fluctuations of laterality suggested flexible recruitment of both hemispheres that is potentially beneficial.

Although larger fluctuations in laterality correlated with better cognitive performance, extreme changes in laterality, i.e., frequent laterality reversal correlated negatively with task performance. Laterality reversal suggests that laterality of a region deviates remarkably from the mean value (the dominant laterality regime, Fig. 1d), i.e., a region’s functional connectivity switches from being more ipsilateral to highly contralateral. Frequent reversal thus indicates that a brain region constantly leaves its dominant functional regime (i.e., more ipsilateral connectivity). Collectively these results suggest that moderate changes in laterality may enhance cognitive performance while extreme laterality changes may hamper cognition. In Results part we have shown that low laterality of the whole brain corresponds to high level of inter-hemispheric communication (or less intra-hemispheric information processing), while high laterality of the whole brain corresponds to the opposite case, these results therefore indicate optimal cognitive performance requires a dynamic balance between inter-hemispheric information exchange and intra-hemispheric information processing. As we also show that low laterality of the whole brain corresponds to low modularity (integration) while high laterality of the whole brain corresponds to high modularity (segregation), our results also echo the recent findings that cognitive function depends on a dynamic, context-sensitive balance between functional integration and segregation ^39^.

In addition to laterality fluctuations and reversal, we furthermore resolved the temporal structures of the whole-brain laterality dynamics. We revealed three recurring states (or ‘meta-states’) with distinct laterality profiles, dwelling time, and transition probabilities. These states generally correspond to the multiple lateralization “axes” identified by Karolis et. al. through dimensionality reduction of 590 meta-analysis maps: our State 1 is characterized by strong left-laterality of the left default mode and language networks, corresponding to the “symbolic communication” axis that involves Broca’s and Wernicke’s areas. Our state 3 demonstrates strong right-laterality of fronto-parietal and default mode networks in the right hemisphere consistent with the “active/perception” axis identified in ^32^.

Laterality of the human brain is hypothesized to be controlled by multiple factors ^9^, which is supported by the distinct spatial clusters identified by the clustering of laterality time series of all brain regions. Three of the four clusters (Cluster 1, 3, 4) we identified roughly correspond to the top three factors identified in ^9^, which include the visual, attention and the default mode network, respectively. It should be noted that our clustering analysis was conducted based on laterality dynamics of each individual, rather than on variations across individuals ^9^.

The four spatial clusters also showed distinct patterns of inter-correlations in their time-varying laterality index (Fig. 3b). Of particular interest is that the time-varying laterality index of Cluster 4 (the default-mode and language network) correlated negatively with those of the other three clusters (Cluster 1, visual network; Cluster 2, sensorimotor network; Cluster 3, attention and FPN network) in most subjects (Cluster 4 and 1, 99.7% subjects with significant negative correlation; Cluster 4 and 2, 99.1%; Cluster 4 and 3, 95.4%). This negative temporal correlation suggests opposite lateralization patterns between Cluster 4 and Cluster 1-3 over different time windows. Furthermore, the more negative the correlation in laterality between Cluster 4 (default-mode and language network) and the other three clusters (especially visual and sensorimotor network), the better the cognitive performance. This suggested that opposite lateralization pattern between Cluster 4 and Cluster 1-3 optimizes parallel processing in the two hemispheres ^14^ and enhances neural capacity. These results could be explained by the causal hypothesis of hemispheric specialization which states that lateralization of one function forces the other function to the opposing hemisphere, which optimize parallel processing in complex tasks and increases processing efficiency ^3,40,41^.

Laterality dynamics of the brain may be related to multiple factors. Functionally, greater lateralization of higher-level brain regions was associated with lower BOLD activity of the whole brain and greater network segregation, suggesting less exchange of information across hemispheres which may be related to less energy (glucose) consumption ^13,42^. Structurally, laterality fluctuations of a region correlated positively with its degree of structural connections. A brain region with wider structural connections may be modulated by multiple systems that likely demonstrate rich patterns of inter-hemisphere interaction, thus larger fluctuations in laterality ^43,44^. Furthermore, we found that pairs of regions with stronger structural connections are more positively correlated in their time-varying laterality index, indicating the role of structural connection in synchronization of functional lateralization of spatially distributed brain regions. In comparison, brain regions with negative inter-correlation of laterality showed less structural connections, possibly affected by more global factors, such as the regulatory effect of neurotransmitters on large-scale brain networks ^39,45,46^.

At the genetic level, structural brain lateralization is known to be heritable in large population samples ^2^. Our research showed that the dynamic characteristics of lateralization were also heritable, especially in those high-order networks such as default-mode and frontal parietal networks, suggesting that laterality dynamics are stable properties of lateralization. From an evolutionary perspective, lateralization arises as a solution to minimize wiring costs while maximizing information processing efficiency in the rapid expansion of the cortex in evolution ^47^. Higher order systems such as default-mode and frontal parietal networks were among the most expanded regions in evolution ^48^. Considering the dynamic laterality of these networks correlate significantly with cognitive performance, the heritability of dynamic lateralization in these networks therefore may confer evolutionary advantages.

## Limitations

In this study, resting-state data was analyzed, so it was not possible to directly relate the results to specific cognitive processes. Future studies should investigate dynamic changes in brain lateralization under task modulation, particularly with multiple task-states or various task loads, to better understand the specific cognitive advantages of dynamic brain lateralization. In addition, age and handedness are important factors potentially affecting lateralization, and handedness is also partly controlled by genetic factors ^49^. Therefore, future studies are warranted that explore how cognitive deterioration with aging may affect lateralization dynamics.

## Conclusion

To conclude, our study demonstrates the dynamic nature of laterality in the human brain at resting-state. We characterized comprehensively the spatiotemporal laterality dynamics by identifying four spatial clusters and three recurring temporal states and demonstrate that the temporal fluctuations of laterality as well as the negative correlation in laterality among different clusters associate with better language task and intellectual performance. We further explored the network-level and structural basis underlying such laterality dynamics, and heritability of laterality dynamics. Our study not only contributes to the understanding of the adaptive nature of human brain laterality in healthy population but may also provide a new perspective for future studies of the genetics of brain laterality and potentially abnormal laterality dynamics in various brain diseases.

## Materials and Methods

### Ethics statement

This paper utilized data collected for the HCP. The scanning protocol, participant recruitment procedures, and informed written consent forms, including consent to share deidentified data, were approved by the Washington University institutional review board ^28^.

#### Dataset

We analyzed multimodal brain imaging data and behavioral measures from the Human Connectome Project 1200 Subjects Release (S1200) ^28^. All brain imaging data were acquired using a multi-band sequence on a 3-Tesla Siemens Skyra scanner. For each participant, four resting-state fMRI scans were acquired: two with right-to-left phase encoding and two with left-to-right phase encoding direction (1200 volumes for each scanning session, TR=0.72 s, voxel size=2×2×2 mm). High-resolution T1-weighted MRI (voxel size= 0.7×0.7×0.7 mm) and diffusion MRI (voxel size=1.25 mm, three shells: b-values=1000, 2000 and 3000 s/mm^2^, 90 diffusion directions per shell) were also acquired for each participant. 991 participants (28.7±3.7 years old, 528 females) were included in the final cohort utilized in this study according to the following criteria: 1) completed all four resting-state fMRI scans; 2) limited in-scanner head motion (mean frame distance (FD) < 0.2 mm); 3) same number of sampling points (i.e., 1200 volumes per scanning session); 4) without any missing data in regions included in the employed parcellation scheme.

#### Resting-state fMRI preprocessing

Resting-state fMRI data were preprocessed using the HCP minimal preprocessing pipeline (*fMRIVolume*) ^50^, and were denoised using the ICA-FIX method ^51^. Then, a spatial smoothing (FHWM=4 mm) and a low-pass filtering (0.01 Hz - 0.1 Hz) were performed. We did not employ global signal regression, as the mean hemispheric time series were used for calculating the lateralization index 11. The whole brain was parcellated into 360 regions (180 for each hemisphere), using the HCP’s multi-modal parcellation (HCP MMP1.0) ^29^. These regions were grouped into 12 functional networks based on the Cole-Anticevic Brain-wide Network Partition (CAB-NP) ^31^.

#### Dynamic laterality Index

The pipeline for the Dynamic Laterality Index (DLI) analysis is illustrated in Fig. 1a. The DLI adopted a sliding-window approach and a traditional laterality index ^11^ to capture the dynamics of laterality. DLI at the tth sliding time window is defined by:

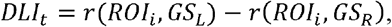

where *ROIi* indicates BOLD time series of ROI *i*, and *GS_L_* and *GS_R_* indicate the global signal, i.e., averaged time series of the voxels within the left and the right hemispheres, respectively. Pearson correlation coefficient *r* is Fisher-z-transformed. There are three_main reasons why we adopted the hemispheric global signals based laterality: 1) Global signal has been shown to be able to reflect and characterize laterality by previous studies 11, and the computational complexity is very low (n*2, n being the number of regions); 2) The whole-brain laterality pattern obtained using hemispheric global signal is highly correlated to that obtained by ROI-based method like autonomy index (AI) defined as follows ^9,10,30,52^:

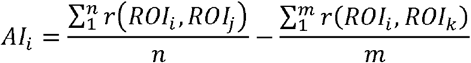

where 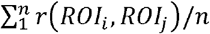 and 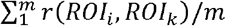 indicate the averaged functional connectivity between BOLD time series of *ROI_i_* and all ROIs in the left and right hemisphere, respectively (Supplemental Fig. 1), and n and m are number of regions in left- and right-hemisphere.

#### Spatial clustering of laterality time-series across the whole-brain

To explore the spatial organization of dynamic lateralization across the whole brain, we performed spatial clustering on the laterality correlation matrix (obtained by calculating Pearson correlation between laterality time series of all pairs of brain regions). Specifically, we applied the Louvain community detection algorithm on the laterality correlation matrix averaged over all 991 subjects and four runs. We set the γ parameter to 1, ran the algorithm for 100 times, and reported the cluster partitioning with the maximum modularity parameter (*Q*).

#### Temporal clustering of whole-brain laterality state

To identify the recurring laterality states of the whole brain from 991 subjects and 1171 time-windows, we adopted a three-stage temporal clustering approach. Firstly, the 1171×4 time-windows of the four runs of each subject were clustered into 10 states by *k*-means clustering. Second, all 9,910 states (991 subjects×10 states) were clustered into 1,000 groups by a second level k-means clustering. Finally, we used Arenas-Fernandez-Gomez (AFG) community detection to cluster the 1000 groups obtained in the second step. AFG allows for multiple resolution screening of the modular structure and a data-driving selection of clustering number. We determined the optimal number of clusters using the modularity coefficients obtained by changing the resolution parameter from 0.1 to 1.5 in steps of 0.1 53. The code of K-means and AFG clustering were from MATLAB function *kmeans* (with the cosine distance metric), and MATLAB Community Detection Toolbox (CDTB v. 0.9, https://www.mathworks.com/matlabcentral/fileexchange/45867-community-detection-toolbox) ^54^, respectively. Based on the final clustering, we took the average pattern of each category as centroids and reclassified all the windows of each subject according to the cosine distance between each window and each centroid.

#### Cognitive performance in behavioral tasks

To understand the cognitive significance of the dynamic brain laterality, we utilized individual performance of a wide variety of cognitive tasks from the NIH Toolbox for Assessment of Neurological and Behavioral function and Penn computerized neurocognitive battery ^55^. These involve story comprehension task that is more left lateralized, Flanker Task (inhibitory control and attention) that maybe more right-lateralized, and Card Sort (cognitive flexibility) and List Sorting task (working memory) that tend to more bilateral. The story comprehension task ^56^ includes a story condition that presents brief auditory stories followed by a binomial forced-choice question, and a math condition to answer addition or subtraction problems. We used three relevant measures: “LanAcc” (“Language_Task_Story_Acc”, the accuracy in answering questions about the story), “LanRT” (“Language_Task_Story_Median_RT”, median response time to answer the questions) and “LanDiff” (“Language_Task_Story_Avg_Difficulty_Level”, the average difficulty of all stories a subject can understand). See Supplemental Method for the details of the measures adopted by other tasks.

### Correlation between dynamic laterality index and cognitive performance

In calculating the correlation between dynamic laterality measures and cognitive performance (unadjusted scores), we used Permutation Analysis of Linear Models (PALM, http://fsl.fmrib.ox.ac.uk/fsl/fslwiki/PALM) to correct for the bias of significance estimation due to the kinship among participants. PALM used exchangeability blocks that is widely adopted to control the family-structure-related bias in HCP data ^57,58^. We calculated the Pearson correlation coefficients between dynamic lateralization attributes [mean laterality index (MLI), laterality fluctuations (LF), laterality reversal (LR) of 4 clusters; laterality correlation within and between 4 clusters; fraction and dwelling time of 3 states, state transitions] and cognitive performances, regressing out sex, age, education years, race, body mass index (BMI), handedness, gray matter volume, white matter volume and head-motion (mean FD). We obtained p values for all correlation coefficients using 5000-times permutation tests. False Discovery Rates (FDR) was used for multiple comparison correction (multiply comparison times=4 MLI + 4 LF + 4 LR + 10 DLI correlation + 3 dwelling time +3 fraction + transition = 29).

#### Association between dynamic laterality, dynamic network properties

To understand the neural and network basis of dynamic brain lateralization, we correlated the laterality time series of each brain region with time-varying brain network measures (which reflect the organization of the brain like modular, or integration property) and the amplitude of low-frequency fluctuation (ALFF) averaged over the whole brain for each subject. We constructed functional brain network using Pearson’s correlation within each sliding time window and computed the following topological measures:

1. Degree centrality (*DC*) ^59,60^, measuring the centrality of each node in the network:

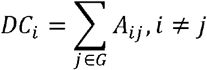 Where *G* is a graph with the node *i* and *j* connected by edge *A_ij_*.
2. Participation coefficient (B^T^) measuring the contribution of each brain region to whole-brain integration:

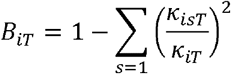 Where *k_isT_* is the strength of the positive connections of region *i* to regions belong to the module *s* at time *T*; *k_iT_* is the sum of strengths of all positive connections of region *i* at time *T*. The participation coefficient of a region is close to 1 if its connections are distributed among all of the modules and 0 if all of its links are within its own module. Community division was obtained by community detection in each time window, and then the participation coefficient was calculated ^22^.
3. Modularity (Q) measuring the tendency of the network to segregate into several independent modules ^61,62^:

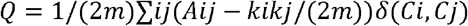 Where *m* is total number of edges in the network; *A* is adjacent matrix; *k_i_* is degree of node *i*; *C_i_* is community assignment of node *i*; when and only when *C_i_*=*C_j_*, *δ(C_i_,C_j_)*=1, otherwise, *δ(C_i_,C_j_)*=0.
4. Inter-hemispheric connection ^63^, measuring the averaged interhemispheric connection strength.

Among these indicators, the averaged *DC* is measured at the global network level, *B_T_* and *Q* are measured at the module level, and interhemispheric connection is measured at the hemispheric level *DC*, *B_T_* and *Q* were calculated using Brain connectivity toolbox (BCT, http://www.brain-connectivity-toolbox.net/) ^64^. We removed all negative connections in line with the general assumptions of graph theory ^60^.

ALFF measures spontaneous energetic activity within each time window ^65^ calculated as follows:

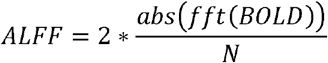

Where *fft* is Fourier transform; *abs* is operator of taking absolute value; *BOLD* is BOLD signal within a time window in any region, and *N* is the length of time window. At global level, we calculated the Pearson *r* between the ALFF/network time series (averaged across the brain) and the absolute laterality time series (averaged across the brain). At local level, we correlated the ALFF/network time series (averaged across the brain) with the absolute laterality time series of each brain region.

#### Structural network analysis

Because the probabilistic fiber tracking is very time-consuming, diffusion tensor imaging (DTI) data of a subset of HCP S1200 release, the “100 Unrelated Subjects” (n=100, 54 females, mean age=29 years) was used for the structural analysis (1 subject was removed due to head motion). Data underwent motion, susceptibility distortion and eddy current distortion correction ^50,66^. We used FMRIB Software Library v6.0 (FSL, https://fsl.fmrib.ox.ac.uk/fsl/fslwiki/) and *MRtrix3*, a toolkit for diffusion-weighted MRI analysis (https://mrtrix.readthedocs.io/en/dev/) ^67^, to construct the structural connectivity (SC) matrix. The processing steps were as follows: 1) tissue segmentation based on T1-weighted structural image; 2) calculation of 4D images with gray matter, white matter and cerebrospinal fluid using multi-shell, multi-tissue constrained spherical deconvolution ^68^; 3) generation of white-matter-constrained tractography using second-order integration over fiber orientation distributions (iFOD2), a probabilistic tracking algorithm (1,000,000 streamlines for each subject, max length=250 mm, cutoff=0.06) ^69^; 4) use of FSL’s FNIRT to reverse-register the MMP parcellation to individual space; 5) calculation of fractional anisotropy (FA) maps using *dtifit* of FSL; 6) measurement of the average FA on all the streamlines connecting any two parcels. This process finally resulted in 360×360 FA matrices and fiber number (FN) matrices for 99 participants. We calculated the FA-degree and FN-degree based on this structural connectivity (SC) matrix, i.e., the sum of FA and FN of each row of the SC matrix. The Spearman correlation between laterality correlation matrix and SC matrix and the correlation between MLI/LF/LR and FN-/FA- degree were calculated for each subject.

#### Heritability analysis

We used twin information in the HCP data to evaluate the heritability of the dynamic laterality measures. In the 991 subjects, 103 pairs of monozygotic twins (MZ), 98 pairs of dizygotic twins (DZ), 195 pairs of sibling (SI) were included, and their zygosity information was used to fit ACE model, which is able to divide total variance in a phenotype (σ^2^) into three additive parts: (A) additive common genetic factor (heritability), (C) common environment, and (E) unshared environment, or other source of measurement error ^70^:

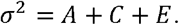

The ratio of the variance explained by A, C and E to the total variance was used as estimates of heritability (*h*^2^), environmental factors (*c*^2^) and error (*e*^2^), all of which sum to 1. All four models (for MLI, LF, LR and laterality correlation matrix) were fitted and estimated using APACE (the Advanced Permutation inference for ACE models, www.warwick.ac.uk/tenichols/apace) ^71^. Before model fitting, sex, age, education years, race, BMI, handedness, gray matter volume, white matter volume and mean FD were regressed out from phenotype as covariables. The significance of heritability for each brain region was obtained using permutation test (1000 permutations per region).

## Code availability

The code used in this study to calculate DLI is available on https://github.com/XinRanWu/Dynamic_Laterality.

## Data Availability

All data files are available from the Human Connectome Project (https://www.humanconnectome.org/study/hcp-young-adult/document/1200-subjects-data-release).

## Author Contributions

X.W, X.K, D.V., and J.Z. contributed to the conception of the study and wrote the paper; X.W. performed the data analyses; P.T. and Z.L. provided critical methodological and conceptual input. B.S., T.W., K.Z. and J.F. provided critical feedback on the manuscript including editing of the manuscript.

## Acknowledgments

Data used in this work were provided by Human Connectome Project (https://www.humanconnectome.org/). JZ was supported by NSFC 61973086, and Shanghai Municipal Science and Technology Major Project (No.2018SHZDZX01) and ZJLab. PMT was supported, in part, by NIH grant U54 EB020403. JF was supported by the 111 Project (No. B18015), the key project of Shanghai Science and Technology (No. 16JC1420402), National Key R&D Program of China (No. 2018YFC1312900), National Natural Science Foundation of China (NSFC 91630314).

