## Supplemental Materials for "Brain Laterality Dynamics Support Human Cognition"

Supplemental Methods

Supplemental Figure 1-7

Supplemental Table 1-3

**Supplemental Methods**

*The relationship between* *dynamic laterality index (DLI) and Autonomy Index (AI).* The traditional laterality index, Autonomy Index ^1-4^ calculates the mean correlation coefficient (functional connectivity) between a ${ROI}_{y}$ and all regions ($x_{i}$) of the left or right hemisphere ($\bar{r}_{xy}=\frac{\sum_{i=1}^{n} r_{x_{i}y}}{n}$) respectively, and used the difference between these two mean correlation coefficients as the measures of ROI’s laterality. In comparison, DLI directly used the correlation coefficients between ${ROI}_{y}$ the global signals of left or right hemisphere ($r_{\bar{x}y}$, where $\bar{x}=\frac{\sum_{i=1}^{n} x_{i}}{n}$) as a substitute, and then calculated the difference between the two correlation coefficients.

We analyze in the following that the two indices are intrinsically related, by showing that the mean correlation coefficient between ${ROI}_{y}$ and regions in one hemisphere is correlated to the correlation coefficient between ${ROI}_{y}$ and the global signal of that hemisphere.

$$\bar{r}_{xy}=\frac{1}{n}\sum_{i=1}^{n} \frac{E\left[ \left( x_{i}-u_{x_{i}} \right)\left( y-u_{y} \right) \right]}{\delta x_{i}\delta y}=\frac{1}{n}\sum_{i=1}^{n} \frac{E\left[ x_{i}y \right]-E\left[ x_{i} \right]E\left[ y \right]}{\delta x_{i}\delta y}$$

$$r_{\bar{x}y}=\frac{E[\left( \bar{x}-u_{\bar{x}} \right)\left( y-u_{y} \right)]}{\delta\bar{x}\delta y}=\frac{E\left[ \bar{x}y \right]-E\left[ \bar{x} \right]E\left[ y \right]}{\sqrt{\delta^{2}\left( \frac{x_{1}+\ldots x_{n}}{n} \right)}\delta y}=\frac{E\left[ \bar{x}y \right]-E\left[ \bar{x} \right]E\left[ y \right]}{\frac{1}{n}\sqrt{\sum_{i=1}^{n} \delta^{2}x_{i}+\sum_{i=1}^{n} \sum_{i\neq j}^{n} Cov\left( x_{i},x_{j} \right)}\delta y}$$

Where $E$ stands for expectation and $\delta$ for standard deviation. It should be noted that only when all $x_{i}$ are independent with each other can $\bar{r}_{xy}$ and $r_{\bar{x}y}$ be highly correlated. For real fMRI data, $x_{i}$s are dependent. Therefore, we also calculated the correlation between AI and DLI based on empirical data to jointly prove the consistency between our indicators and previous indicators (see Methods).

*The HCP language task.* The language task was developed by Binder et al. ^5^. In the story blocks, participants were presented with brief auditory stories adapted from Aesop’s fables, followed by a 2-alternative forced-choice question to check the participants’ understanding of the story topic. For example, after a story about an eagle that saves a man who had done him a favor, participants were asked, “Was that about revenge or reciprocity?”. In the math blocks, participants were presented auditory series of addition and subtraction (e.g., “fourteen plus twelve”), followed by “equals” and then two choices (e.g., “twenty-nine or twenty-six”). To ensure similar level of difficulty across participants, math trials automatically adapted to the participants responses. The story and math trials were matched in terms of auditory, duration, attention demand and phonological input.

**Supplemental Figures**

***Supplemental Figure 1****. Repeatability of 3 dynamic laterality measures across sessions* *(A) Left: the average Dynamic Lateralization Index (DLI) map of four sessions of HCP data. Right: correlation coefficient among the four sessions (average among all the subjects). (B) Left: the average Autonomy Index (AI) map calculated by four sessions of HCP data. Right: correlation coefficient between the four sessions (average among all the subjects). It can be seen that DLI shows higher inter-session repeatability than AI. (C) Repeatability of various dynamic laterality measures across different window length. REST1/REST2 indicates the scan time (in first or second day). LR/RL indicates the scanning direction. LR, left to right. RL, right to left.*

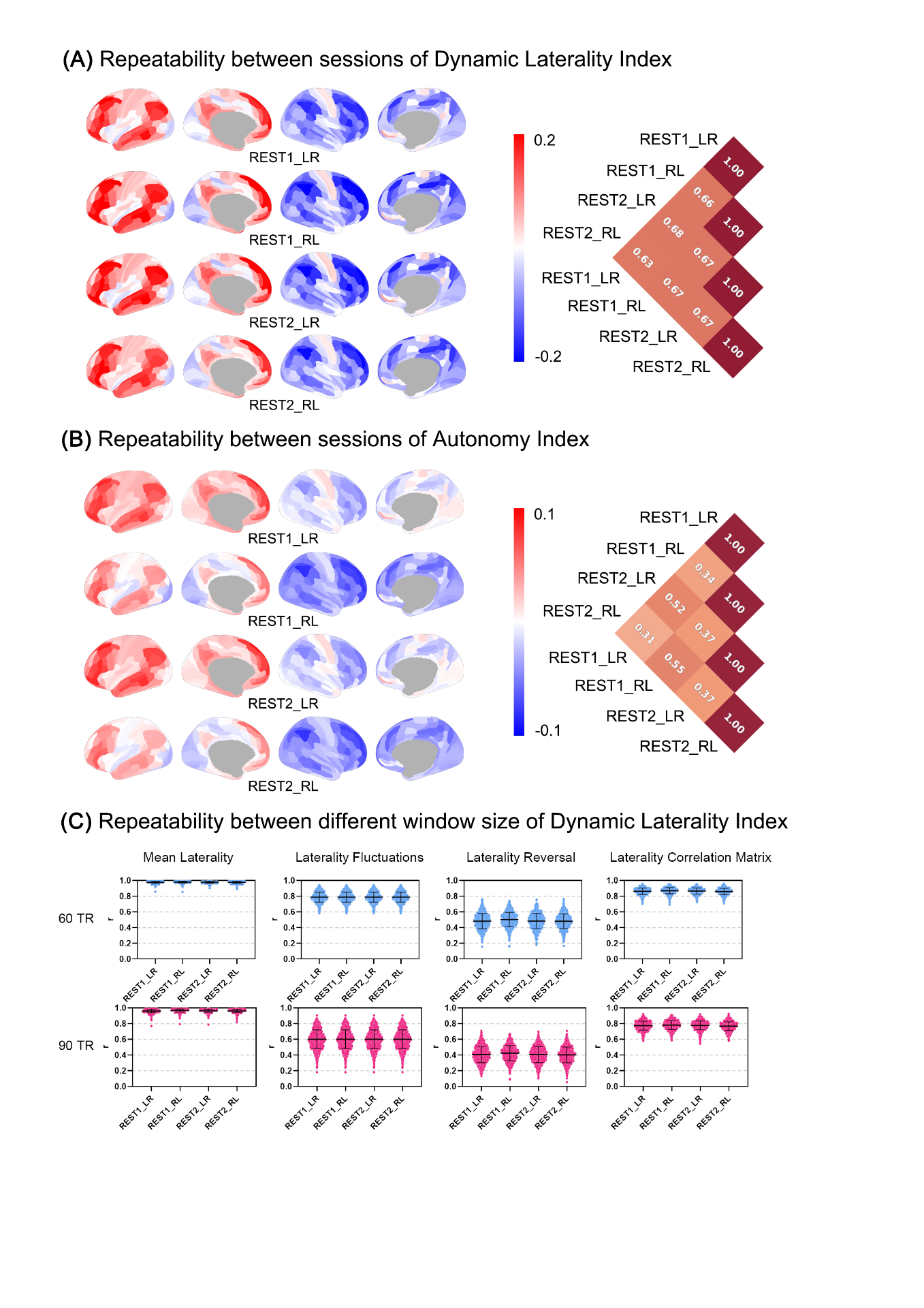

***Supplemental Figure 2.*** *Average homotopic laterality correlation.*

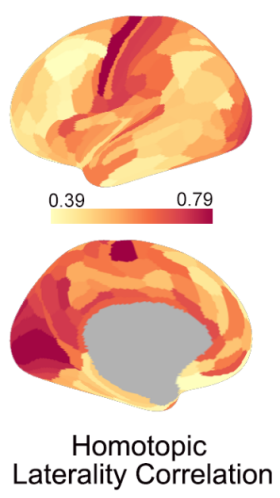

***Supplemental Figure 3.*** *(A) Dynamic laterality measures (mean laterality index, laterality fluctuations and laterality reversal) across four clusters. The statistical test was repeated-measure ANOVA, with the asterisk representing significance (***, p<0.01, ****, <0.001) (B) The average laterality correlation matrix over all 991 subjects (arranged by 12 functional networks of the Cole-Anticevic Brain-wide Network Partition).*

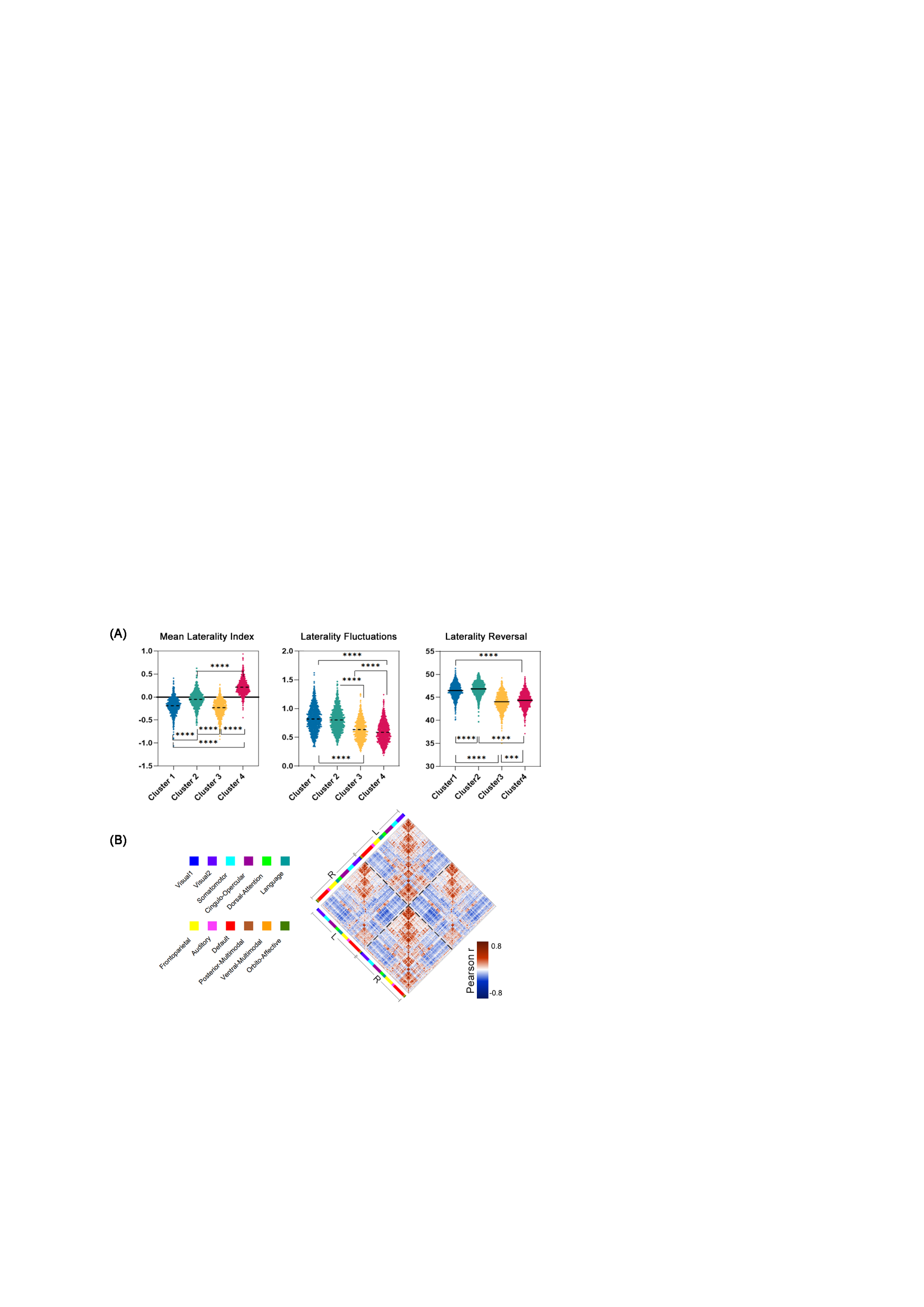

***Supplemental Figure 4.*** *(A-E) The association between the dynamic laterality index with the average time series of network graph theory indicators and Amplitude of Low Frequency Fluctuations (ALFF) of each brain region. The values in the brain maps represent the t-value of the Pearson correlation coefficient r (one sample t-test).*

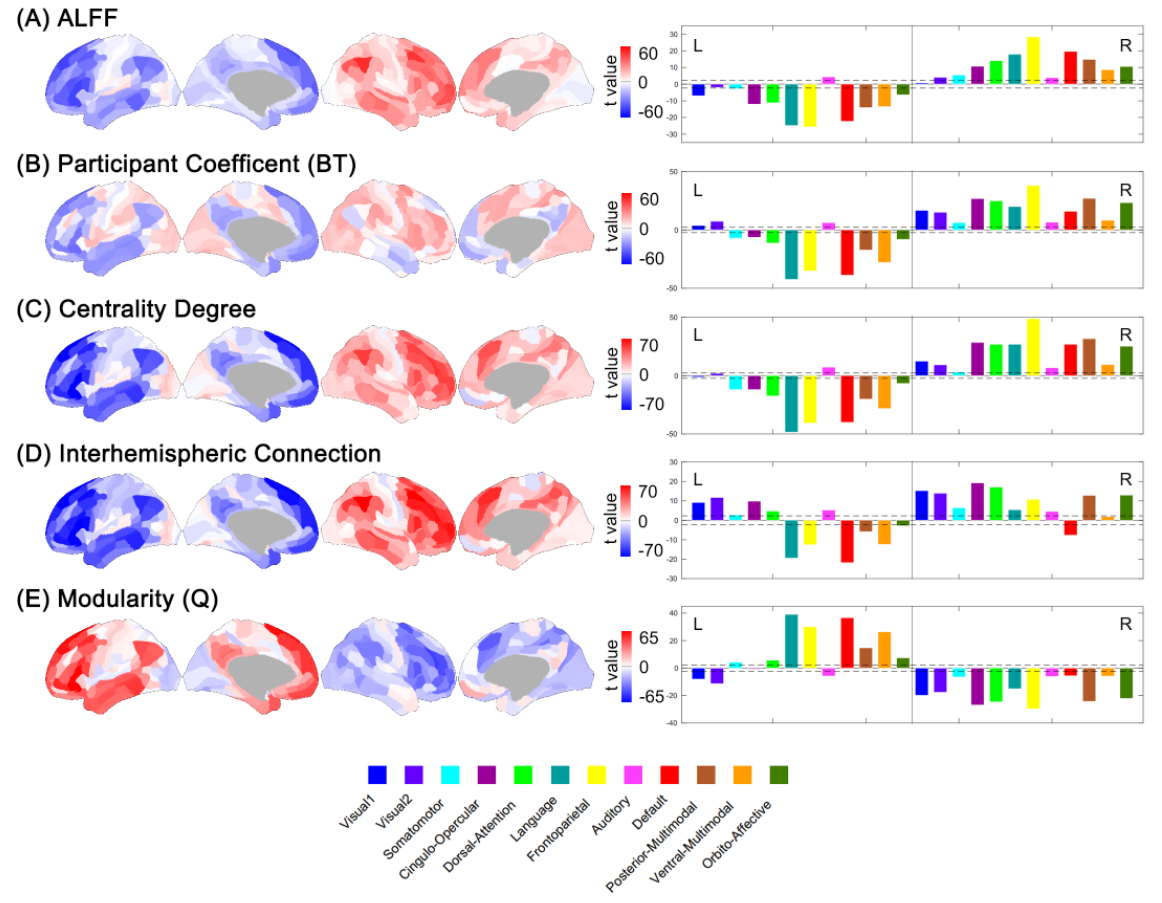

***Supplemental Figure 5.*** *The statistical differences of lateralization correlation matrix, ALFF and network index between every pair of the three temporal states. The paired t test was used, and all values shown in the figure were t statistics. Only regions with t value exceeded the significant threshold (±2.33, p<0.001) were shown in brain maps.*

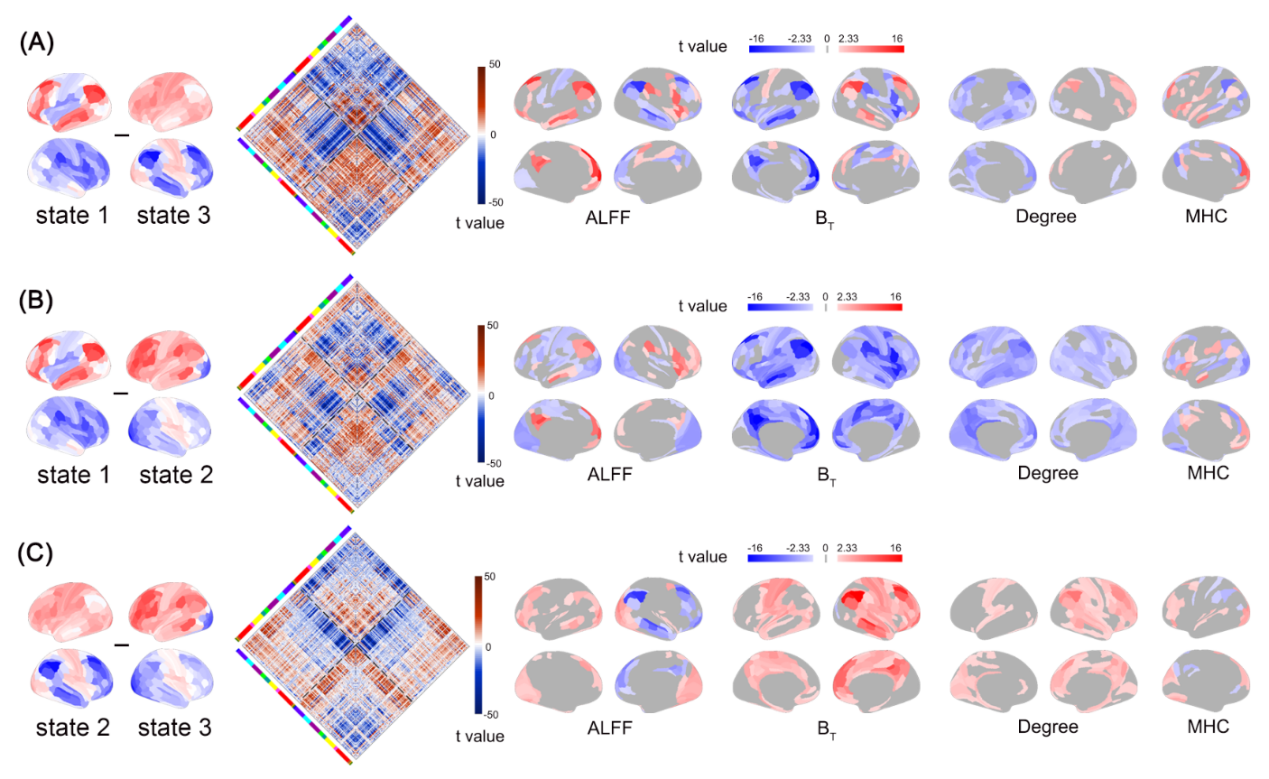

*Supplemental Figure 6. The statistical differences of Q between every pair of the three temporal states. The one-way ANOVA and post-hoc test were used.*

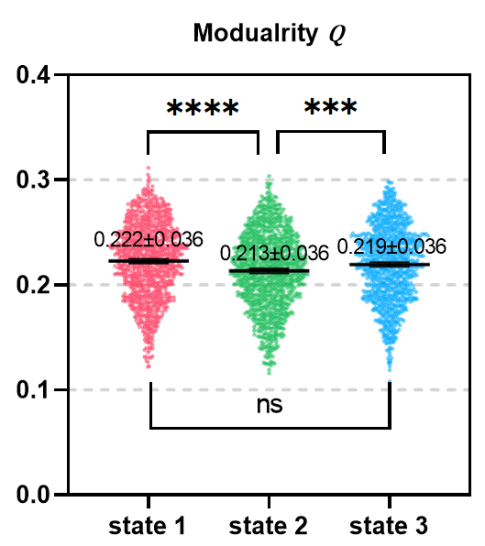

*Supplemental Figure 7. (A) Mean structural connection matrix. (B) Mean fiber number (FN) degree. (C) fractional anisotropy (FA) degree. (D) The distribution of Spearman correlation coefficients between structural connection attributes [FN-degree, FA-degree and structural connection matrix (SCM, based on FN or FA)] and dynamic laterality measures [LF: laterality fluctuations, LR: laterality reversal and Laterality Correlation Matrix (LCM)].*

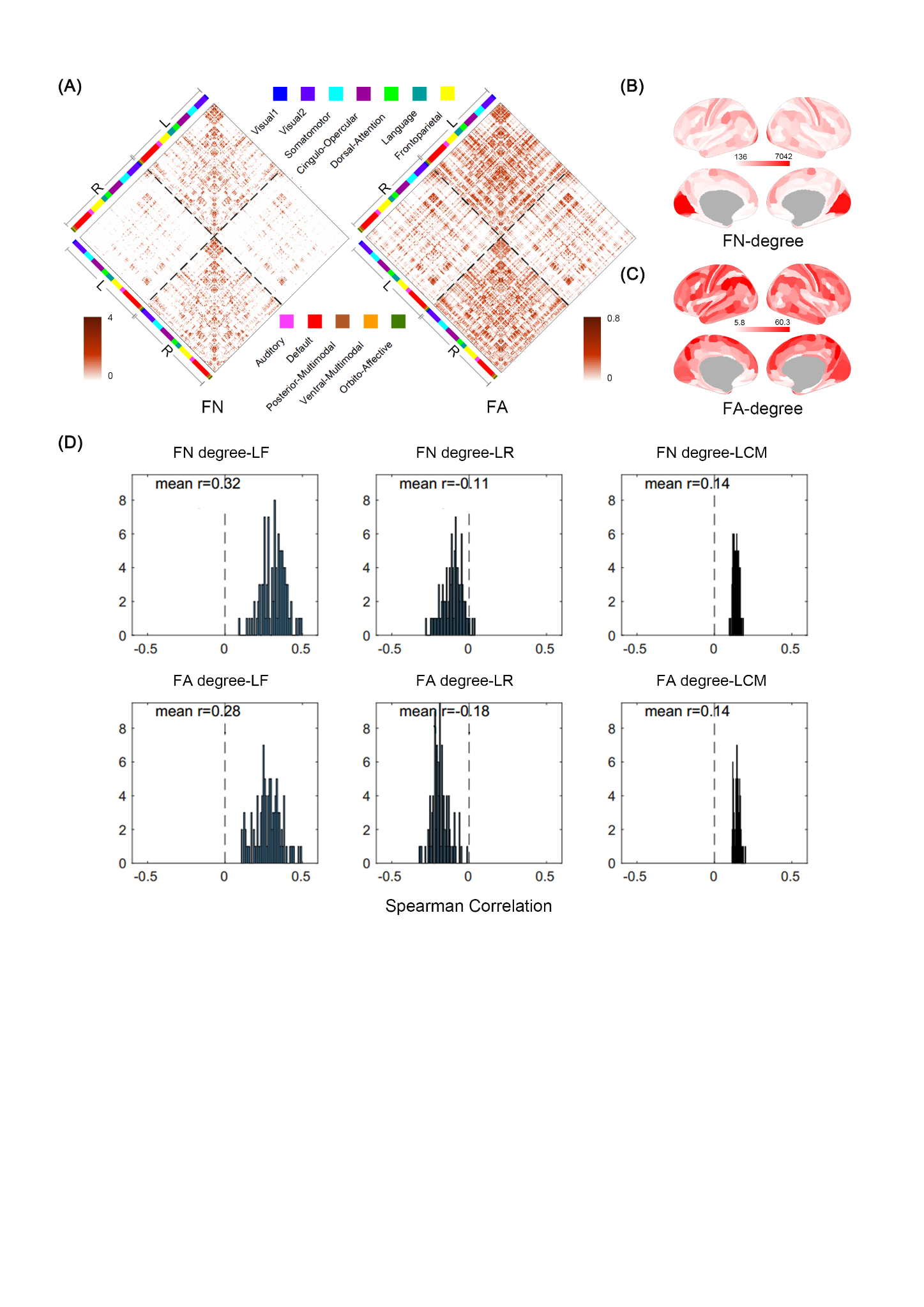

***Supplemental Figure 8.*** *The association between the averaged time series of absolute value of dynamic laterality index (abs(DLI)) with the average time series of network graph theory indicators and Amplitude of Low Frequency Fluctuations (ALFF).* *We selected data from three subjects, sub-100206, sub-100307, and sub-100408, to show relationship between global lateralization (represented by averaged absolute value of DLI) and dynamic ALFF/ graph theory indicators. The images show that higher global lateralization is associated with lower ALFF and stronger whole-brain dissociation (higher modularization Q, lower participant coefficient, centrality degree, and interhemispheric connectivity).*

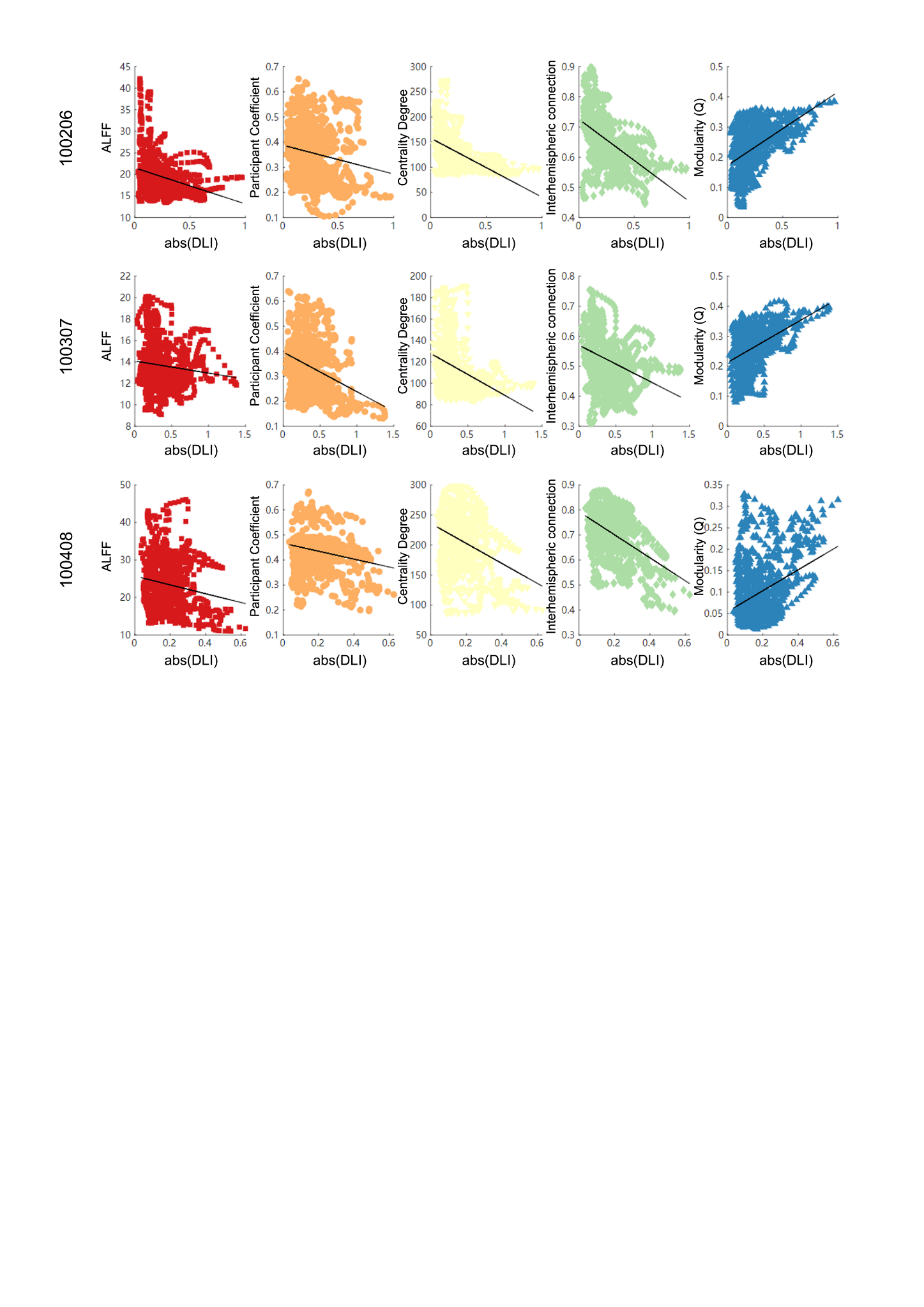

**Supplemental Tables**

***Supplemental Table 1.*** *The association between dynamic lateralization characteristic of four clusters and cognitive abilities. Each row of the table corresponds to a cognitive test score in the HCP. PicVocab, Picture Vocabulary Test;* *ReadEng, Reading Test (reading decoding skill); Cardsort, Dimensional Change Card Sort Test (executive function, specifically tapping cognitive flexibility); Flanker, Flanker task (executive function, specifically tapping inhibitory control and attention); ProcSpeed, Pattern Comparison Processing Test (speed of processing); PicSeq, Picture Sequence Memory Test (episodic memory); VSPLOT_TC, Total Number Correct of Penn Line Orientation. PMAT24_A_CR, Number of Correct Responses of Penn Matrix Test (non-verbal reasoning);* *ListSort,* *List Sorting Working Memory Test; IWRD_TOT, Total Number of Correct Responses of Penn Word Memory; LanAcc, the accuracy of answering questions about the story of language task; LanRT, median response time to answer the questions of language task; LanDiff, the average difficulty of all stories of language task for each subject which represents the overall language comprehension ability. r, Pearson correlation coefficient; p, the permutation test significance obtained by Permutation Analysis of Linear Models (PALM). The bold text represents the association remains significant after False Discovery Rates (FDR) correction (q =0.05). See HCP Data Dictionary for more details.*

|  |  | ***Mean Laterality Index*** | | | | ***Laterality Fluctuations*** | | | | ***Laterality Reversal*** | | | |
| --- | --- | --- | --- | --- | --- | --- | --- | --- | --- | --- | --- | --- | --- |
|  |  | ***Cluster 1*** | ***Cluster 2*** | ***Cluster 3*** | ***Cluster 4*** | ***Cluster 1*** | ***Cluster 2*** | ***Cluster 3*** | ***Cluster 4*** | ***Cluster 1*** | ***Cluster 2*** | ***Cluster 3*** | ***Cluster 4*** |
| ***PicVocab*** | ***r*** | 0.04 | -0.02 | -0.11 | 0.05 | 0.07 | 0.08 | 0.09 | 0.05 | 0.04 | -0.02 | -0.12 | -0.08 |
|  | ***p*** | 0.358 | 0.540 | 0.033 | 0.365 | 0.227 | 0.112 | 0.060 | 0.350 | 0.335 | 0.634 | 0.022 | 0.149 |
| ***ReadEng*** | ***r*** | 0.03 | -0.06 | -0.04 | 0.03 | 0.08 | 0.11 | 0.08 | 0.07 | 0.02 | -0.01 | -0.11 | -0.10 |
|  | ***p*** | 0.384 | 0.109 | 0.467 | 0.566 | 0.130 | 0.015 | 0.091 | 0.151 | 0.700 | 0.733 | 0.044 | 0.058 |
| ***CardSort*** | ***r*** | -0.05 | -0.08 | -0.14 | 0.13 | 0.12 | 0.14 | 0.14 | 0.14 | -0.05 | -0.06 | -0.15 | -0.15 |
|  | ***p*** | 0.118 | 0.025 | **0.001** | **0.001** | **0.001** | **0.000** | **0.000** | **0.000** | 0.190 | 0.079 | **0.000** | **0.000** |
| ***Flanker*** | ***r*** | 0.02 | -0.01 | -0.09 | 0.05 | 0.06 | 0.05 | 0.07 | 0.05 | 0.01 | -0.01 | -0.10 | -0.09 |
|  | ***p*** | 0.630 | 0.856 | 0.018 | 0.160 | 0.085 | 0.165 | 0.084 | 0.164 | 0.889 | 0.725 | 0.016 | 0.029 |
| ***ProcSpeed*** | ***r*** | -0.02 | -0.05 | -0.10 | 0.10 | 0.07 | 0.06 | 0.09 | 0.09 | -0.08 | -0.05 | -0.14 | -0.12 |
|  | ***p*** | 0.638 | 0.141 | **0.009** | **0.005** | 0.075 | 0.113 | **0.009** | 0.021 | 0.045 | 0.131 | **0.001** | **0.003** |
| ***VSPLOT_TC*** | ***r*** | 0.01 | -0.05 | -0.08 | 0.06 | 0.06 | 0.06 | 0.06 | 0.04 | 0.01 | 0.03 | -0.08 | -0.05 |
|  | ***p*** | 0.718 | 0.173 | 0.056 | 0.093 | 0.188 | 0.130 | 0.133 | 0.292 | 0.835 | 0.387 | 0.071 | 0.317 |
| ***PMAT24_A_CR*** | ***r*** | 0.01 | -0.01 | -0.08 | 0.05 | 0.04 | 0.05 | 0.06 | 0.01 | -0.02 | -0.04 | -0.11 | -0.07 |
|  | ***p*** | 0.693 | 0.770 | 0.067 | 0.288 | 0.396 | 0.323 | 0.201 | 0.874 | 0.513 | 0.228 | 0.012 | 0.102 |
| ***PicSeq*** | ***r*** | -0.04 | 0.01 | -0.04 | 0.06 | 0.04 | 0.02 | 0.01 | 0.02 | 0.02 | -0.02 | -0.06 | -0.03 |
|  | ***p*** | 0.278 | 0.701 | 0.236 | 0.065 | 0.256 | 0.673 | 0.745 | 0.598 | 0.597 | 0.617 | 0.113 | 0.368 |
| ***ListSort*** | ***r*** | 0.02 | 0.00 | -0.04 | 0.03 | 0.00 | 0.00 | -0.02 | -0.03 | 0.02 | -0.01 | -0.09 | -0.03 |
|  | ***p*** | 0.495 | 0.979 | 0.365 | 0.429 | 0.952 | 0.913 | 0.724 | 0.528 | 0.654 | 0.699 | 0.039 | 0.495 |
| ***IWRD_TOT*** | ***r*** | -0.01 | 0.00 | -0.06 | 0.04 | 0.05 | 0.05 | 0.06 | 0.05 | 0.00 | -0.04 | -0.06 | -0.04 |
|  | ***p*** | 0.792 | 0.922 | 0.056 | 0.200 | 0.169 | 0.134 | 0.065 | 0.147 | 0.963 | 0.288 | 0.054 | 0.293 |
| ***LanAcc*** | ***r*** | 0.03 | -0.03 | -0.04 | 0.01 | 0.01 | 0.02 | 0.02 | 0.02 | -0.04 | -0.07 | -0.11 | -0.05 |
|  | ***p*** | 0.298 | 0.430 | 0.267 | 0.674 | 0.730 | 0.562 | 0.545 | 0.678 | 0.306 | 0.038 | 0.004 | 0.145 |
| ***LanRT*** | ***r*** | 0.08 | 0.04 | 0.01 | -0.05 | -0.07 | -0.07 | -0.05 | -0.08 | 0.10 | 0.00 | 0.08 | 0.08 |
|  | ***p*** | 0.018 | 0.254 | 0.830 | 0.129 | 0.074 | 0.073 | 0.163 | 0.042 | 0.007 | 0.970 | 0.042 | 0.047 |
| ***LanDiff*** | ***r*** | -0.01 | -0.06 | -0.12 | 0.11 | 0.11 | 0.16 | 0.11 | 0.11 | -0.05 | -0.03 | -0.20 | -0.14 |
|  | ***p*** | 0.836 | 0.133 | **0.002** | **0.010** | 0.033 | **0.000** | **0.014** | **0.018** | 0.293 | 0.302 | **0.000** | **0.002** |

***Supplemental Table 2.*** *The association between Laterality Covariance among four clusters and cognitive abilities. Each row of the table corresponds to a cognitive test score in the HCP. r, Pearson correlation coefficient; p, the permutation test significance obtained by PALM. The bold text represents the association remains significant after False Discovery Rates (FDR) correction (q =0.05).*

|  |  | ***Laterality Covariance*** | | | | | | | | | |
| --- | --- | --- | --- | --- | --- | --- | --- | --- | --- | --- | --- |
|  |  | ***Cluster 1-1*** | ***Cluster 1-2*** | ***Cluster 1-3*** | ***Cluster 1-4*** | ***Cluster 2-2*** | ***Cluster 2-3*** | ***Cluster 2-4*** | ***Cluster 3-3*** | ***Cluster 3-4*** | ***Cluster 4-4*** |
| ***PicVocab*** | ***r*** | 0.02 | 0.05 | -0.06 | 0.04 | 0.02 | 0.01 | -0.04 | 0.05 | 0.00 | -0.01 |
|  | ***p*** | 0.723 | 0.309 | 0.125 | 0.326 | 0.597 | 0.817 | 0.419 | 0.179 | 0.914 | 0.781 |
| ***ReadEng*** | ***r*** | 0.02 | 0.09 | -0.02 | 0.03 | 0.08 | 0.03 | -0.08 | -0.01 | -0.01 | 0.00 |
|  | ***p*** | 0.707 | 0.055 | 0.685 | 0.496 | 0.072 | 0.512 | 0.054 | 0.734 | 0.889 | 0.926 |
| ***CardSort*** | ***r*** | 0.07 | 0.06 | 0.10 | -0.04 | 0.05 | 0.05 | -0.09 | 0.06 | -0.12 | 0.11 |
|  | ***p*** | 0.052 | 0.098 | **0.011** | 0.211 | 0.155 | 0.103 | **0.012** | 0.093 | **0.001** | **0.003** |
| ***Flanker*** | ***r*** | 0.05 | 0.03 | 0.03 | -0.01 | -0.01 | 0.04 | -0.02 | 0.04 | -0.08 | 0.04 |
|  | ***p*** | 0.103 | 0.373 | 0.498 | 0.830 | 0.723 | 0.186 | 0.542 | 0.231 | 0.034 | 0.301 |
| ***ProcSpeed*** | ***r*** | 0.01 | 0.08 | 0.05 | -0.02 | -0.02 | 0.02 | -0.05 | 0.04 | -0.10 | 0.06 |
|  | ***p*** | 0.808 | 0.039 | 0.251 | 0.629 | 0.509 | 0.480 | 0.152 | 0.172 | **0.009** | 0.081 |
| ***VSPLOT_TC*** | ***r*** | 0.04 | 0.05 | -0.04 | 0.01 | 0.04 | 0.01 | -0.03 | 0.04 | -0.01 | 0.03 |
|  | ***p*** | 0.397 | 0.275 | 0.245 | 0.743 | 0.395 | 0.744 | 0.534 | 0.202 | 0.726 | 0.543 |
| ***PMAT24_A_CR*** | ***r*** | 0.07 | -0.03 | -0.07 | 0.04 | 0.07 | 0.00 | 0.03 | 0.09 | 0.02 | -0.01 |
|  | ***p*** | 0.098 | 0.519 | 0.078 | 0.255 | 0.086 | 0.993 | 0.407 | 0.014 | 0.725 | 0.860 |
| ***PicSeq*** | ***r*** | 0.06 | 0.04 | -0.01 | 0.00 | 0.01 | 0.01 | -0.01 | 0.00 | -0.03 | 0.02 |
|  | ***p*** | 0.129 | 0.374 | 0.879 | 0.992 | 0.822 | 0.839 | 0.678 | 0.914 | 0.505 | 0.547 |
| ***ListSort*** | ***r*** | 0.03 | -0.03 | -0.03 | 0.04 | 0.02 | -0.05 | 0.04 | -0.01 | 0.04 | -0.04 |
|  | ***p*** | 0.449 | 0.528 | 0.381 | 0.211 | 0.624 | 0.125 | 0.210 | 0.882 | 0.365 | 0.279 |
| ***IWRD_TOT*** | ***r*** | 0.03 | 0.05 | 0.00 | 0.00 | 0.04 | 0.00 | -0.05 | 0.05 | -0.06 | 0.04 |
|  | ***p*** | 0.369 | 0.138 | 0.947 | 0.957 | 0.242 | 0.989 | 0.164 | 0.099 | 0.055 | 0.302 |
| ***LanAcc*** | ***r*** | 0.01 | 0.04 | -0.02 | 0.03 | 0.02 | 0.02 | -0.02 | 0.03 | 0.00 | -0.01 |
|  | ***p*** | 0.843 | 0.231 | 0.587 | 0.427 | 0.648 | 0.500 | 0.571 | 0.332 | 0.990 | 0.743 |
| ***LanRT*** | ***r*** | -0.01 | -0.04 | -0.04 | 0.04 | -0.01 | -0.03 | 0.04 | 0.08 | 0.01 | -0.03 |
|  | ***p*** | 0.703 | 0.234 | 0.217 | 0.189 | 0.721 | 0.290 | 0.253 | 0.021 | 0.731 | 0.428 |
| ***LanDiff*** | ***r*** | 0.03 | 0.11 | 0.01 | 0.00 | 0.11 | 0.02 | -0.09 | 0.03 | -0.07 | 0.06 |
|  | ***p*** | 0.519 | **0.016** | 0.777 | 0.985 | **0.011** | 0.564 | 0.033 | 0.453 | 0.097 | 0.173 |

***Supplemental Table 3.*** *The association between characteristics of three laterality states and cognitive abilities. Each row of the table corresponds to a cognitive test score in the HCP. r, Pearson correlation coefficient; p, the permutation test significance obtained by PALM. The bold text represents the association remains significant after False Discovery Rates (FDR) correction (q =0.05).*

|  |  | ***State Fraction*** | | | ***State Mean Dwelling Time*** | | | ***Transition*** |
| --- | --- | --- | --- | --- | --- | --- | --- | --- |
|  |  | ***State 1*** | ***State 2*** | ***State 3*** | ***State 1*** | ***State 2*** | ***State 3*** |  |
| ***PicVocab*** | ***r*** | -0.01 | -0.06 | 0.07 | 0.01 | -0.04 | 0.08 | -0.04 |
|  | ***p*** | 0.808 | 0.071 | 0.038 | 0.729 | 0.257 | 0.034 | 0.329 |
| ***ReadEng*** | ***r*** | 0.01 | -0.09 | 0.07 | 0.03 | -0.08 | 0.05 | 0.01 |
|  | ***p*** | 0.723 | 0.015 | 0.045 | 0.440 | 0.020 | 0.133 | 0.912 |
| ***CardSort*** | ***r*** | 0.10 | -0.08 | -0.03 | 0.10 | -0.05 | 0.01 | -0.09 |
|  | ***p*** | **0.006** | **0.014** | 0.428 | **0.004** | 0.134 | 0.829 | **0.009** |
| ***Flanker*** | ***r*** | 0.03 | -0.05 | 0.01 | 0.02 | -0.02 | 0.02 | -0.03 |
|  | ***p*** | 0.422 | 0.164 | 0.698 | 0.542 | 0.544 | 0.551 | 0.357 |
| ***ProcSpeed*** | ***r*** | 0.09 | -0.09 | -0.01 | 0.06 | -0.07 | 0.01 | -0.02 |
|  | ***p*** | 0.017 | **0.010** | 0.785 | 0.069 | 0.044 | 0.847 | 0.554 |
| ***VSPLOT_TC*** | ***r*** | 0.03 | -0.06 | 0.02 | 0.00 | -0.06 | -0.02 | 0.06 |
|  | ***p*** | 0.305 | 0.085 | 0.598 | 0.883 | 0.077 | 0.643 | 0.111 |
| ***PMAT24_A_CR*** | ***r*** | -0.01 | 0.00 | 0.01 | 0.00 | 0.01 | 0.01 | 0.00 |
|  | ***p*** | 0.706 | 0.969 | 0.661 | 0.901 | 0.842 | 0.675 | 0.952 |
| ***PicSeq*** | ***r*** | 0.01 | 0.04 | -0.04 | -0.02 | 0.00 | -0.07 | 0.08 |
|  | ***p*** | 0.811 | 0.319 | 0.267 | 0.603 | 0.986 | 0.078 | 0.022 |
| ***ListSort*** | ***r*** | 0.03 | -0.02 | -0.01 | 0.03 | -0.03 | -0.03 | 0.03 |
|  | ***p*** | 0.418 | 0.627 | 0.699 | 0.444 | 0.391 | 0.318 | 0.363 |
| ***IWRD_TOT*** | ***r*** | 0.01 | -0.02 | 0.01 | 0.02 | -0.03 | -0.01 | 0.01 |
|  | ***p*** | 0.888 | 0.548 | 0.714 | 0.587 | 0.409 | 0.861 | 0.821 |
| ***LanAcc*** | ***r*** | 0.01 | -0.06 | 0.04 | 0.01 | -0.04 | 0.04 | -0.01 |
|  | ***p*** | 0.772 | 0.096 | 0.232 | 0.769 | 0.224 | 0.297 | 0.715 |
| ***LanRT*** | ***r*** | -0.03 | 0.04 | 0.00 | -0.04 | 0.03 | -0.02 | 0.03 |
|  | ***p*** | 0.361 | 0.270 | 0.970 | 0.273 | 0.424 | 0.474 | 0.282 |
| ***LanDiff*** | ***r*** | 0.06 | -0.10 | 0.03 | 0.07 | -0.09 | 0.02 | -0.02 |
|  | ***p*** | 0.076 | **0.007** | 0.368 | 0.041 | **0.015** | 0.625 | 0.611 |
